# *Veillonella dispar* triggers “dual-hits” in the pathogenesis of autoimmune hepatitis and primary biliary cholangitis via testosterone depletion-BAFF-B cell axis and Th17 cell activation

**DOI:** 10.64898/2026.09.26.754610

**Authors:** Ya-Ling Liu, Qing-Qing Xing, Xiang Lin, Wei-Wei Zeng, Yin-Hong Chen, Yan-Ying You, Li-Na Zhou, Xiao-Ning Chen, Min-Xia Wu, Han-Xin Xue, Shangeng Weng, Jin-Shui Pan

**Author notes:** Correspondence: Dr. Jin-Shui Pan, Department of Hepatology, Hepatology Research Institute, the First Affiliated Hospital of Fujian Medical University, 20th, Chazhong Road, Fuzhou, Fujian 350005, China., Dr. Shangeng Weng, Department of Hepatobiliary and Pancreatic Surgery, Hepatology Research Institute, the First Affiliated Hospital of Fujian Medical University, 20th, Chazhong Road, Fuzhou, Fujian 350005, China. These authors share first authorship (Ya-Ling Liu, Qing-Qing Xing, Xiang Lin, Wei-Wei Zeng).

## Abstract

**Objective:** Autoimmune hepatitis (AIH) and primary biliary cholangitis (PBC) frequently present as overlap syndromes, characterized by shared sexual dimorphism and elevated intestinal *Veillonella dispar* (*V. dispar*) abundance, yet exhibiting distinct hallmark pathological manifestations. This study aimed to elucidate the shared pathogenesis, phenotypic differentiation mechanisms and sex-specific predisposition of AIH and PBC induced by *V. dispar*.

**Methods:** Clean-grade and germ-free (GF) mice were employed to establish AIH and PBC models. Model induction involved oral gavage of *V. dispar* and intraperitoneal injection of either liver-specific antigen S100 (for AIH) or a pyruvate dehydrogenase complex E2 subunit (PDC-E2) mimetic peptide (for PBC). Single-cell RNA sequencing, metabolomics, transmission electron microscopy, western blotting, and flow cytometry were utilized to elucidate the underlying mechanisms.

**Results:** *V. dispar* successfully induced autoimmune liver disease-like features in mice, characterized by significant intrahepatic B cell infiltration. Mechanistically, *V. dispar* caused systemic arginine reduction, leading to intrinsic apoptosis of gonadal stromal cells and a subsequent reduction in testosterone. This decreased testosterone weakened its inhibitory effect on B cell activating factor (BAFF) secretion, thereby amplifying B cell-mediated immune responses. Furthermore, in the presence of S100 or PDC-E2 mimetic peptide, *V. dispar* also facilitated Th17 cell activation, which guided the phenotypic differentiation into AIH or PBC, respectively. Consistent with these findings, our clinical observations revealed that patients with AIH and PBC tended to harbor enriched *V. dispar*, along with lower serum arginine and testosterone. In addition, elevated serum levels of BAFF and IL-17 were also observed in these patients.

**Conclusion:** *V. dispar* induces reduced testosterone levels and upregulation of BAFF, leading to intrahepatic B cell expansion, which serves as the common pathogenic basis for AIH and PBC. The presence of S100 and PDC-E2 subsequently drives further maturation of these expanded B cells and promotes Th17 cell activation, thereby determining the phenotypic divergence between AIH and PBC.

## 1. Introduction

Autoimmune hepatitis (AIH) and primary biliary cholangitis (PBC) are two prevalent autoimmune liver diseases (AILD) characterized by chronic, progressive inflammation that can lead to cirrhosis and liver failure, often necessitating liver transplantation. These conditions are globally distributed, affecting individuals across all races and ethnicities, and have shown an increasing prevalence in recent years [1–4]. For nearly five decades, corticosteroids and azathioprine have been the largely unchanged first-line regimen for AIH [5, 6]. Ursodeoxycholic acid (UDCA) is the first-line therapy for PBC [7, 8]. While UDCA can alleviate symptoms and delay disease progression in some patients, it is not a curative treatment. Consequently, a significant number of patients respond poorly to UDCA. For these individuals, or those experiencing end-stage PBC or severe intractable pruritus, liver transplantation remains the only viable treatment option [7–9]. However, even after transplantation, PBC can recur, with rates as high as 30%–35% [10, 11]. The mechanisms of AIH and PBC involve environmental triggers, immunogenetic susceptibility, innate and adaptive immune abnormalities, gut microbiota dysbiosis, and sex hormone levels [12–14]. The gut-liver axis links gut microbiota dysbiosis to abnormalities in immune tolerance, an early and pivotal event in the onset of ALD [15, 16]. Recent studies have demonstrated that the abundance of intestinal *Veillonella dispar* (*V. dispar*) is positively correlated with serum aspartate aminotransferase (AST) levels and intrahepatic inflammation. Furthermore, gut *V. dispar* abundance significantly decreased in PBC patients after UDCA treatment, but remained elevated in non-responders [17, 18], indicating its association with UDCA responsiveness. Similar to that, the patients with AIH also have enriched intestinal *V. dispar* [19].

Both AIH and PBC demonstrate a significant female predominance in incidence. For AIH, the female-to-male ratio typically ranges from approximately 4:1 to 6:1, whereas for PBC, this ratio extends from 4:1 to 9:1 [13, 20]. Similarly, other autoimmune diseases also show a female predominance, such as Sjögren’s syndrome (10:1 female-to-male ratio), systemic lupus erythematosus (SLE), systemic sclerosis, and antiphospholipid syndrome (all with approximately 5:1 female-to-male ratios [21]). Collectively, these findings suggest that sex hormones may critically modulate immune tolerance and inflammatory responses.

AIH and PBC both show intestinal *V. dispar* enrichment and female predominance. Furthermore, sequential or simultaneous PBC-AIH overlap syndrome may develop [22, 23]. Thus, a shared pathogenic mechanism between PBC and AIH is plausible. However, significant differences exist between PBC and AIH, including clinical presentation, autoantibody profiles, and histopathological changes. Among susceptible females with enriched *V. dispar*, some develop AIH while others manifest PBC, suggesting a disease-specific determining factor beyond the shared mechanism. Moreover, the intrinsic association between *V. dispar* enrichment, sexual dimorphism, and the proliferation and activation of B and T cells remains unclear. Therefore, in-depth research in this area is is crucial to elucidate the key pathogenic mechanisms of AIH and PBC.

## 2. Materials and methods

### 2.1 Animal models

Female and male C57BL/6 mice (6-8 weeks old) were acquired from GemPharmatech Co., Ltd. (Jiangsu, China). Germ-free C57BL/6J mice were also purchased from the same supplier. All mice were housed in a clean-grade animal facility under controlled conditions with adequate ventilation. Germ-free C57BL/6J mice were maintained under standardised conditions, including a 12-hour light/12-hour dark cycle, a temperature of 22°C±2°C, and ad libitum access to food and water. All mice were randomly assigned to the experimental and control groups, and no statistically significant differences in baseline parameters were observed between the groups.

C57BL/6 mice were euthanized to extract the liver antigen S100 [24]. Mice in the model group received an intraperitoneal injection of S100 protein (0.25 mg/250uL) fully emulsified in an equivalent volume of complete Freund’s adjuvant (CFA; Sigma-Aldrich, St. Louis, MO, USA) on days 0 and 7. Control mice received an intraperitoneal injection of 250 uL of phosphate-buffered saline emulsified with an equivalent volume of complete Freund’s adjuvant.

The PDC-E2 163-176 peptide (-GDLLAEIETDKATI-) is an immunodominant autoreactive T-cell epitope in patients with PBC [25]. Mimicry peptides were synthesized and purified by Sangon Biotech Co., Ltd. (Shanghai, China). The peptide was dissolved in chilled phosphate-buffered saline at 20 mg/mL and subsequently diluted as required. Mice in the model group received intraperitoneal injections of an emulsified mixture of peptide and CFA (100ug/50ul) on days 0, 7, and 14. Ctrl group mice were intraperitoneally injected with an equivalent volume of the vehicle at identical time points. All experimental procedures were conducted in accordance with guidelines approved by the Institutional Animal Care and Use Committee of Fujian Medical University (IACUC FJMU 2025-Y-1047).

## 3. Results

### 3.1 *V. dispar* induced AIH/PBC-like histological changes

To explore the pathogenic role of *V. dispar* in murine liver, mice were randomly assigned to two groups and administered either PBS or *V. dispar* every other day for 20 weeks (Fig. S1A). Experimental data revealed that oral gavage with *V. dispar* significantly increased serum biochemical parameters, including AST, ALP, TC, and TG, in both male and female mice (Fig. 1A). Furthermore, significant lymphocyte infiltration and an elevated histological activity index (HAI) were observed in the portal areas of both male and female mice (Fig. 1C, E), suggesting a role for *V. dispar* in the initiation and progression of AIH and PBC-like histological manifestions. To further validate the involvement of *V. dispar* in inducing AIH and PBC-like histological changes, germ-free mice were colonized with *V. dispar*. This colonization resulted in similar elevations in specific biochemical parameters (Fig. 1B) and portal inflammation, consistent with observations in clean-grade mice (Fig. 1D, F).

**Figure 1.**
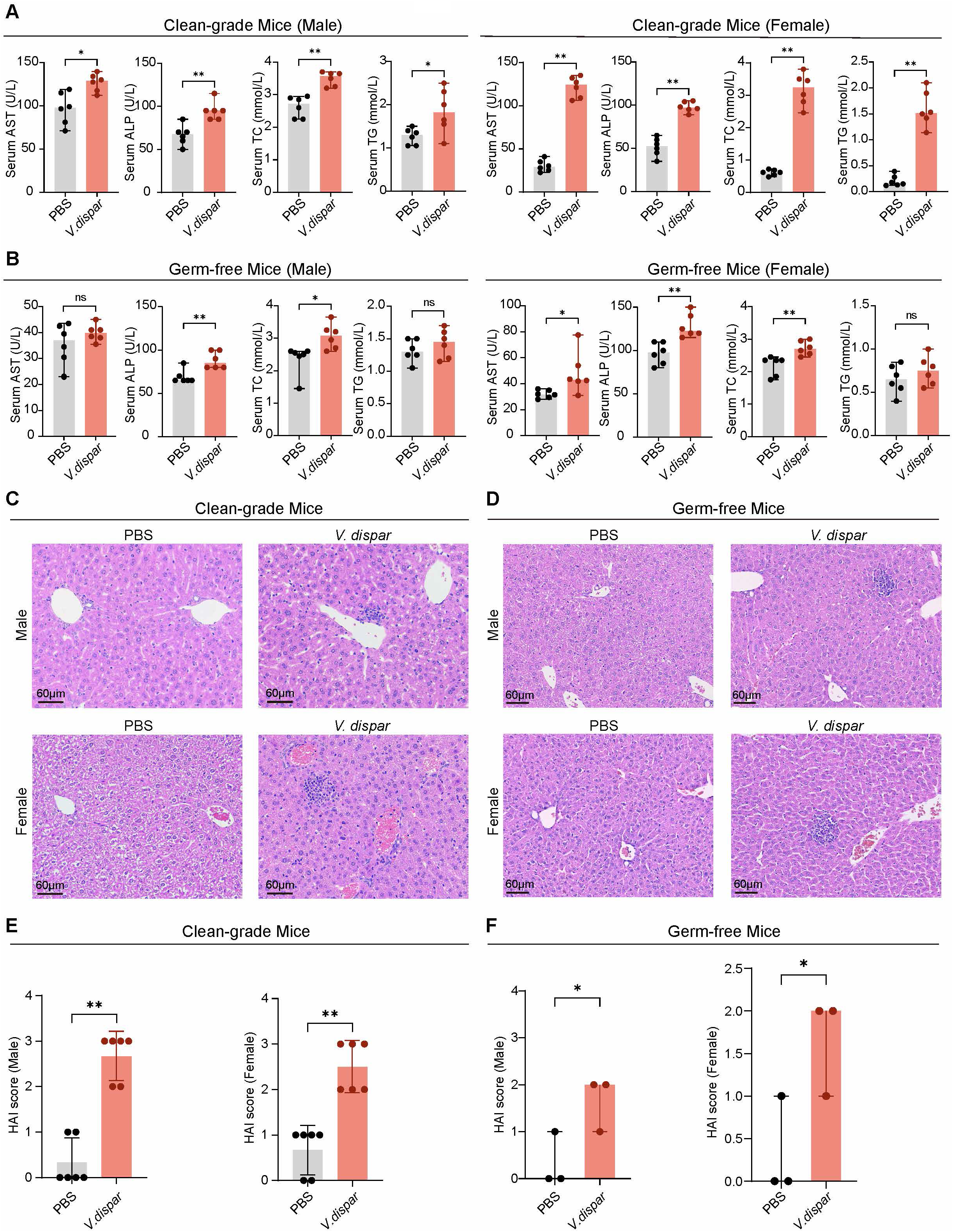
*V. dispar* induces AIH/PBC-like liver histopathological alterations. (A, B) Serum levels of aspartate aminotransferase (AST), alkaline phosphatase (ALP), total cholesterol (TC), and triglyceride (TG) in male and female clean-grade mice (A) and germ-free (GF) mice (B). Mice were orally gavaged with PBS or *V. dispar* every 2 days for 90 days. Serum biochemistry was analysed to evaluate hepatic inflammation. *V. dispar* significantly elevated AST, ALP, TC, and TG levels, indicating hepatic inflammation (n = 6). (C, D) Representative H&E-stained liver sections from clean-grade mice (C) and GF mice (D). (E, F) Liver injury was evaluated based on the histological activity index (HAI). *V. dispar* exacerbated portal inflammation in both sexes (n = 3–6). Data are presented as median with 95% confidence interval. \**P* < 0.05, \*\**P* < 0.01, \*\*\**P* < 0.001; ns, not significant. Data shown in (A), (B), and (E) were conducted using the Mann-Whitney U test.

### 3.2 *V. dispar* promoted intrahepatic B-cell infiltration

C57BL/6 mice were gavaged with PBS or *V. dispar* for 12 weeks, and liver tissues were then subjected to scRNA-seq (Fig. 2A). This analysis yielded 357,513 and 353,048 liver cells from the PBS and *V. dispar* groups, respectively. Dimensionality reduction and clustering identified 36 subpopulations across 11 major hepatic cell lineages (Fig. 2B, C). Compared to controls, scRNA-seq revealed a significant increase in hepatic B cells in the *V. dispar* group, while the proportions of T cells, natural killer (NK) cells, and monocytes remained unchanged (Fig. 2D). These findings were consistently validated across multiple experimental approaches. Flow cytometry confirmed a significant increase in intrahepatic B cell infiltration in both sexes after *V. dispar* gavage, whereas total CD3^+^ T cells and their related subsets showed no appreciable changes among the groups (Fig. 2E, F and S1B). Additionally, spleen flow cytometric analysis demonstrated a marked elevation in splenic B cells in both sexes, whereas no substantial differences were observed in total CD3^+^ T cells or their major subsets among the groups (Fig. S1C, D and E). Immunofluorescence staining further confirmed the infiltration of CD19^+^ B cells in the liver and spleen of clean-grade male mice, paralleling the flow cytometric results from both the liver and spleen (Fig. S1F). To assess the role of the microbiota, male germ-free mice were similarly gavaged for 12 weeks. Immunofluorescence staining in these mice revealed an increase in CD19+ B cells in both the liver (Fig. 1G) and spleen (Fig. 1H), while the proportions of CD3^+^ cells, CD3^+^CD4^+^ cells, and CD3^+^CD8^+^ cells in the spleen remained unchanged (Fig. 1I, J).

**Figure 2.**
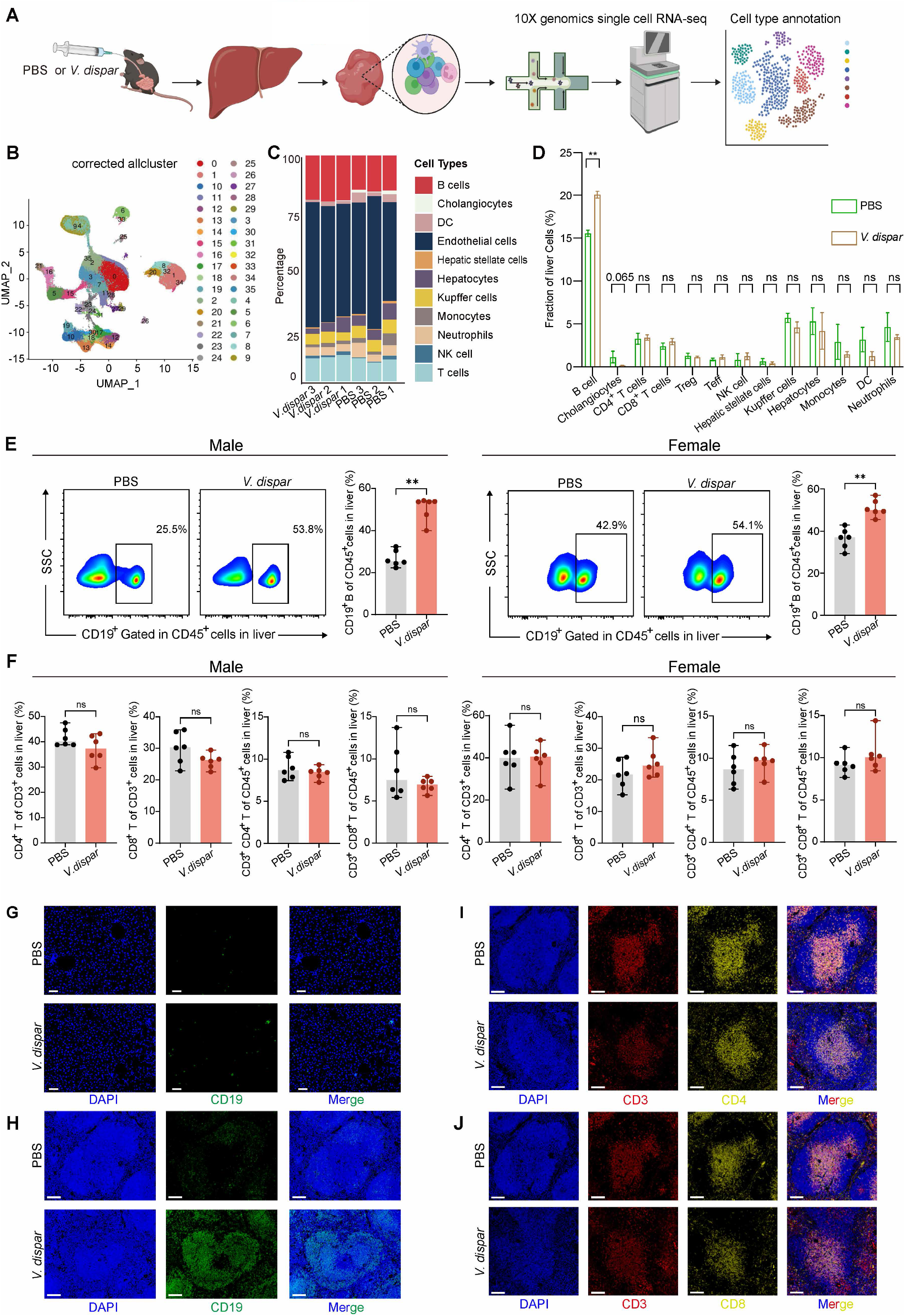
*V. dispar* drives intrahepatic B cell infiltration in mice. (A) C57BL/6 mice were orally gavaged with PBS or *V. dispar* for 12 weeks, followed by 10x Genomics single-cell RNA sequencing (scRNA-seq) of liver tissues. (B, C) UMAP visualization (B) and cellular composition analysis (C) identified 36 cell clusters encompassing 11 major intrahepatic cell lineages. (D) The *V. dispar*-treated group demonstrating a strong increase in intrahepatic B cells and a reduction in cholangiocytes, with no significant changes in hepatocytes, T cells, or NK cells. (E) Representative flow cytometric plots showing elevated frequencies of CD19^+^ B cells (gated on CD45^+^ leukocytes) in male and female livers following *V. dispar* administration (n = 6). (F) No significant differences in hepatic CD4^+^ or CD8^+^ T cell frequencies were detected between groups in either sex (n = 6). (G, H) Representative immunofluorescence images of CD19 (green) indicating B cell infiltration in the liver (G) and spleen (H). (I, J) Representative immunofluorescence images of the spleen showing DAPI (nuclear staining), CD3, and CD4; DAPI, CD3, and CD8. Data are presented as median with 95% confidence interval. \**P* < 0.05, \*\**P* < 0.01, \*\*\**P* < 0.001; ns, not significant. Statistical analyses for (E, F) were performed using the Mann-Whitney U test; for (D) using the Kruskal-Wallis test with Dunn’s multiple comparisons correction.

### 3.3 *V. dispar* upregulated BAFF expression

Compared with the controls, BAFF levels in both serum and liver were significantly increased in the *V. dispar*-treated mice (Fig. 3A, B). Correlation analysis revealed a strong positive correlation between serum and liver BAFF levels and intrahepatic B cell infiltration in both sexes (Fig. 3C, D), suggesting that BAFF may mediate B cell proliferation and recruitment. To further investigate this mechanism, pseudo-germ-free mice were generated by orally administering multiple antibiotics for one week, followed gavage with PBS or *V. dispar*. Subsequently, these mice were treated with anti-BAFF or an isotype control (Fig. 3E). Immunofluorescence staining demonstrated that intrahepatic BAFF levels were elevated by *V. dispar* gavage + isotype treatment, and reduced by *V. dispar* gavage + anti-BAFF treatment (Fig. S2A). Immunofluorescence and H&E staining revealed that *V. dispar* gavage increased intrahepatic B cell infiltration, which was subsequently hindered by anti-BAFF treatment (Fig. 3F and S2B). Flow cytometry revealed that serum BAFF levels were elevated by *V. dispar* gavage and isotype treatment but reduced by *V. dispar* gavage and anti-BAFF treatment, with intrahepatic B cell infiltration demonstrating a comparable change in both sexes (Fig. 3G, H).

**Figure 3.**
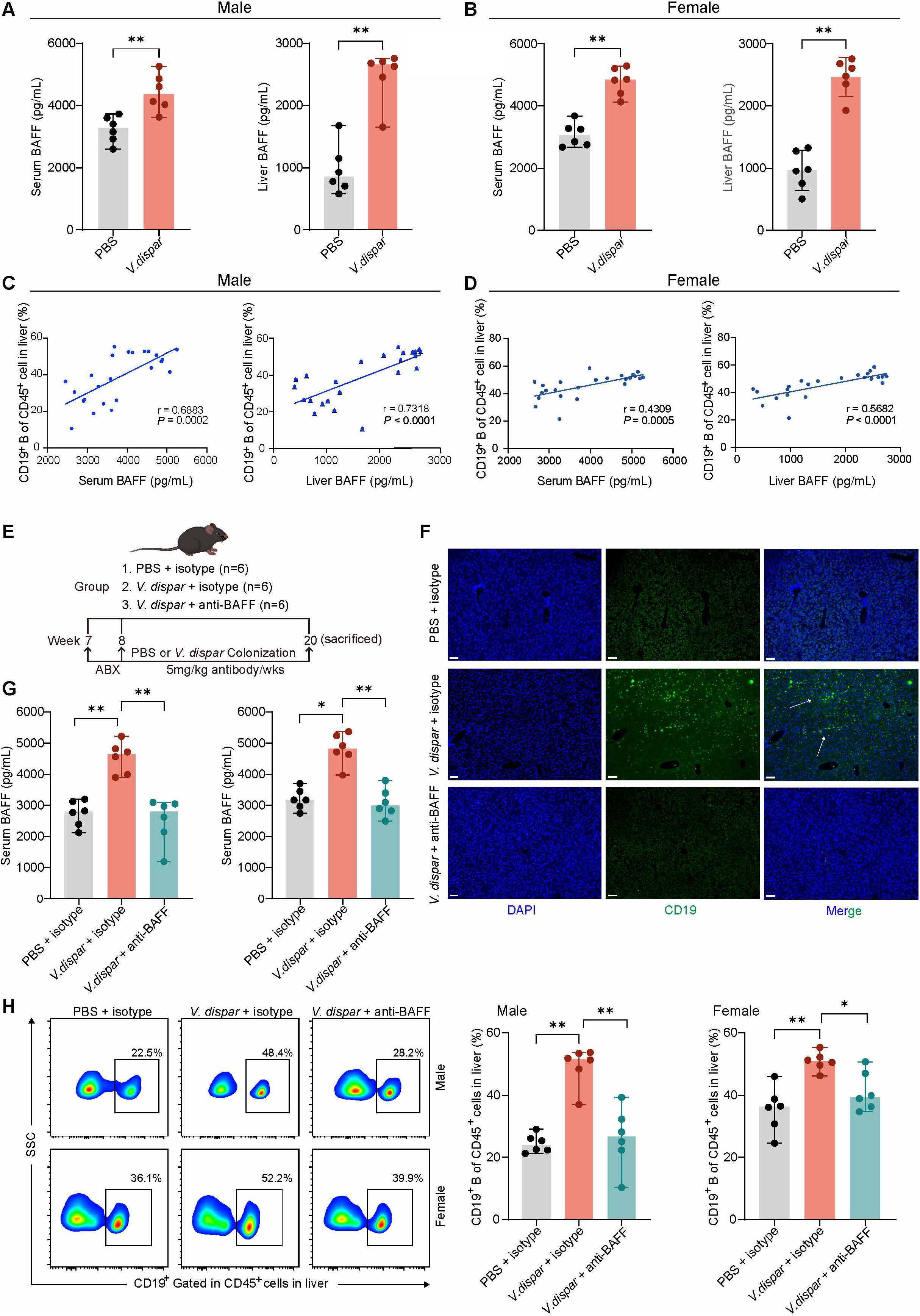
*V. dispar* upregulates BAFF expression to promote intrahepatic B cell infiltration. (A, B) ELISA quantification of serum and hepatic BAFF levels in male (A) and female (B) mice. BAFF concentrations were significantly higher in *V. dispar*-treated mice (n = 6). (C, D) Spearman’s rank correlation between serum/hepatic BAFF levels and hepatic CD19^+^ B cell frequencies in male (C) and female (D) mice. (E) Experimental design: antibiotic-pretreated mice were gavaged with *V. dispar* or PBS and treated weekly with anti-BAFF or isotype-control antibodies. (F) Representative immunofluorescence staining of CD19 (green) showing intrahepatic B cell accumulation induced by *V. dispar* and attenuated by anti-BAFF treatment. Scale bar: 100 μm. (G) Serum BAFF levels (ELISA) were elevated in the *V. dispar* + isotype group and reduced by anti-BAFF in both sexes (n = 6). (H) Representative flow cytometric plots and quantification demonstrating increased intrahepatic CD19^+^ B cell frequencies induced by *V. dispar* and their reduction by anti-BAFF in both sexes (n = 6). Data are presented as median with 95% confidence interval. \**P* < 0.05, \*\**P* < 0.01, \*\*\**P* < 0.001; ns, not significant. Statistical analyses for (A, B) were conducted using the Mann-Whitney U test; for (C, D) using Spearman’s rank correlation; for (G, H) using the Kruskal-Wallis test with Dunn’s multiple comparisons correction.

### 3.4 *V. dispar* decreased serum testosterone levels and conversely elevated BAFF secretion

Serum hormone levels were assessed in male and female mice 12 weeks after gavage with PBS or *V. dispar*. *V. dispar* treatment significantly reduced serum testosterone levels in both sexes (Fig. 4A, B and C). Previous studies indicate that testosterone suppresses the proliferation of splenic fibroblastic reticular cells (FRCs) and their BAFF production [26]. To validate this finding, 6–8-week-old C57BL/6 mice were divided into the following groups: sham operation + *V. dispar* (Sham + *V. dispar*), gonadectomy + *V. dispar* (ORX + *V. dispar*), and gonadectomy + *V. dispar* + testosterone supplementation (ORX + *V. dispar* + T) (Fig. 4D). H&E staining and flow cytometry revealed that testosterone supplementation reduced the elevated intrahepatic B-cell infiltration in ORX mice (Fig. 4E, G). B cells infiltration in the spleen was comparable to that in the liver (Fig. S3C, D). The ORX + *V. dispar* group showed a significant decrease in serum testosterone levels, which was subsequently restored in the ORX + *V. dispar* + T group (Fig. S3A, B). In both sexes, serum BAFF levels were significantly elevated following gonadectomy + *V. dispar* treatment, an effect that was subsequently reversed by gonadectomy + *V. dispar* + testosterone supplementation (Fig. 4F).

**Figure 4.**
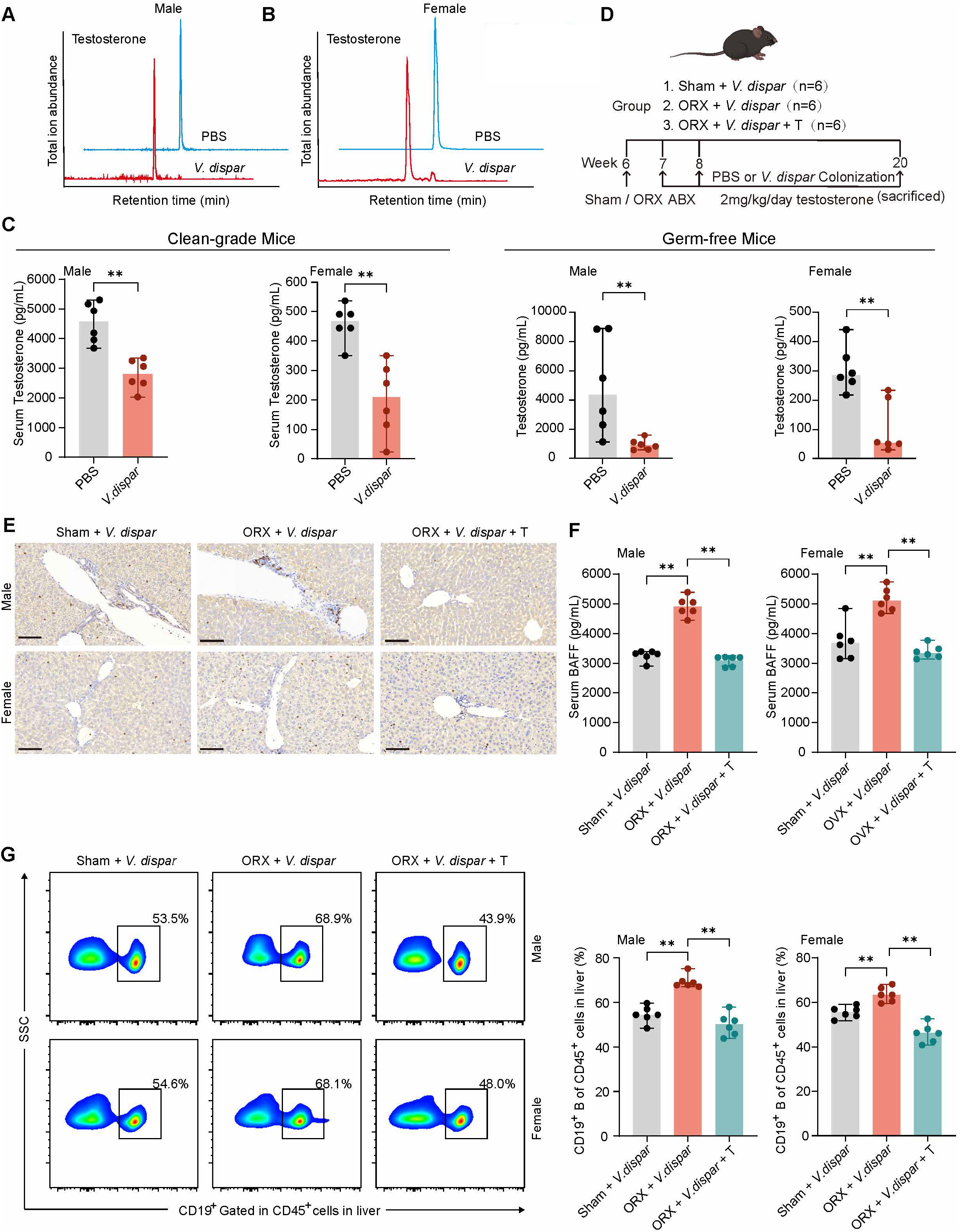
*V. dispar* reduces testosterone levels to enhance BAFF expression and B cell infiltration. (A–C) Representative LC-MS/MS spectra (A, B) and quantification (C) of serum testosterone in male and female mice. *V. dispar* significantly lowered testosterone levels in both sexes (n = 6). (D) Experimental design: Mice underwent sham or orchiectomy (ORX) surgery with or without testosterone (T) supplementation, followed by *V. dispar* gavage. (E) Representative immunohistochemical staining of liver sections shows increased CD19^+^ B cells in ORX mice, which was mitigated by testosterone supplementation (n = 6). Scale bar: 100 μm. (F) Serum BAFF levels were elevated in ORX + *V. dispar* mice and reduced by testosterone supplementation (n = 6). (G) Representative flow cytometric plots show increased hepatic CD19^+^ B cells in ORX + *V. dispar* mice, which was reversed by testosterone supplementation in both sexes (n = 6). Data are presented as median with 95% confidence interval. \**P* < 0.05, \*\**P* < 0.01, \*\*\**P* < 0.001; ns, not significant. Data in (A) and (C) were analyzed using the Mann-Whitney U test; for (F, G) using the Kruskal-Wallis test with Dunn’s multiple comparisons correction.

### 3.5 *V. dispar* induced intrinsic apoptosis of gonadal stromal cells

Gonadal stromal cells are the primary source of testosterone in both male and female mice. To investigate the effects of *V. dispar* on gonadal function, we performed histological and molecular analyses of these tissues. Following 12 weeks of *V. dispar* administration by gavage, no significant effect on the testis-to-body weight ratio was observed in clean-grade mice (Fig. S4A). However, both clean-grade and germ-free mice exhibited loosening of the testicular and ovarian stroma. Histological examination of the testes further revealed a decreased number and disorganized arrangement of germ cells, along with structural disintegration between the seminiferous tubules and the interstitial space (Fig. 5A and Fig. S4B). To assess apoptosis, immunofluorescence staining for Caspase-3 in ovaries and 3β-HSD and TUNEL assays in testes confirmed a significant increase in apoptotic cells following *V. dispar* treatment (Fig. 5B, C, and Fig. S4C).

**Figure 5.**
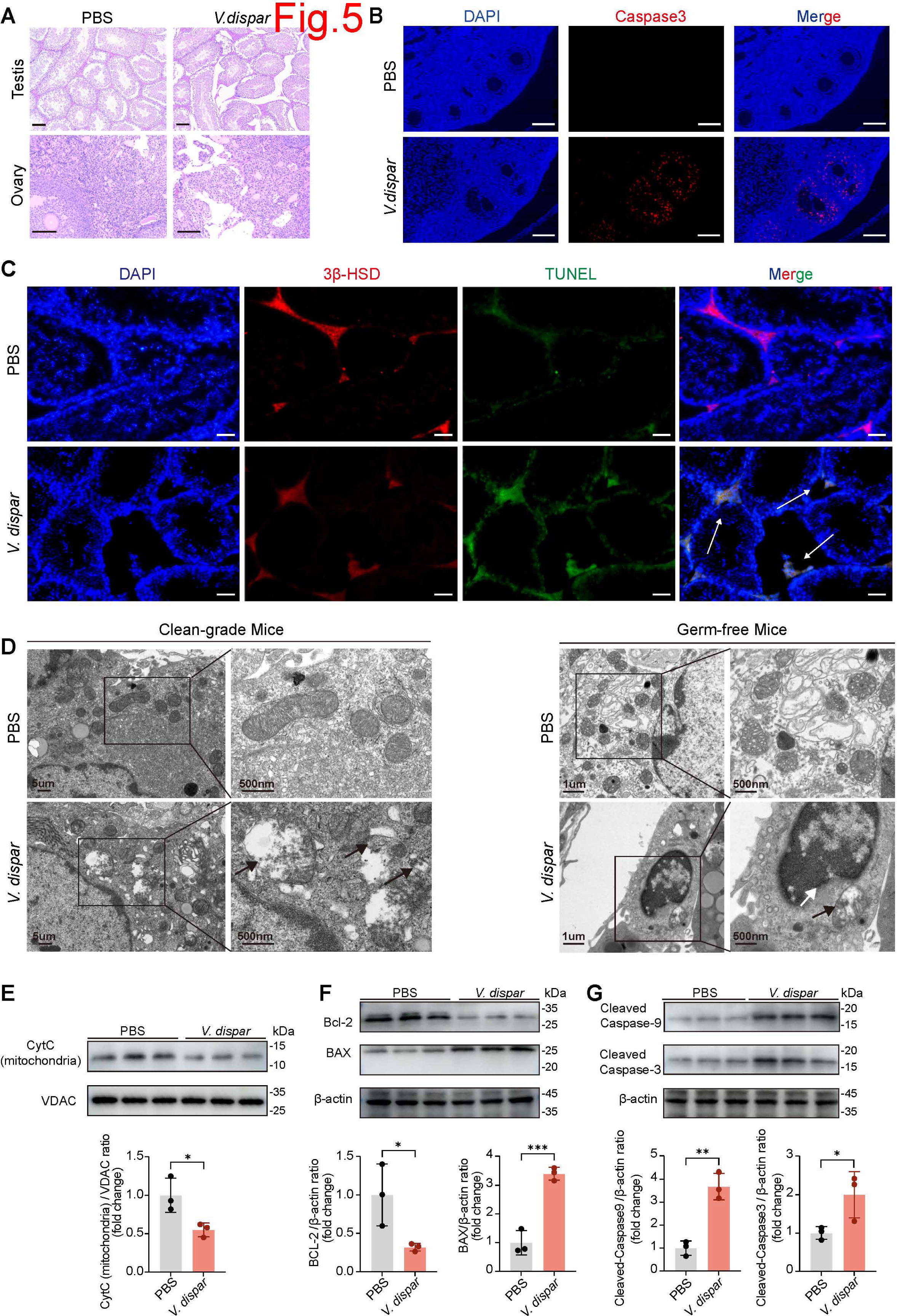
*V. dispar* triggers intrinsic apoptosis in gonadal stromal cells. (A) Representative H&E staining of testes and ovaries shows reduced germ cells, loosened stroma, and disorganized interstitium after *V. dispar* treatment. Scale bar: 100 μm. (B) Representative immunofluorescence of cleaved Caspase-3 (green) in ovaries shows increased apoptotic cells following *V. dispar* administration. DAPI (blue) labels nuclei. Scale bar: 100 μm. (C) Representative immunofluorescence of 3β-HSD (red), TUNEL (green), and DAPI (blue) in testes shows stromal apoptosis and loss of 3β-HSD^+^ cells in *V. dispar*-treated mice. Scale bar: 20 μm. (D) Transmission electron microscopy images show mitochondrial damage and apoptotic body formation in testicular stromal cells after *V. dispar* treatment. (E) Western blotting shows decreased mitochondrial cytochrome C (CytC) in testes of *V. dispar*-treated mice (n = 3). (F) Expression of pro- and anti-apoptotic proteins in primary testicular stromal cells, showing increased Bax and decreased Bcl-2 (n = 3). (G) Elevated cleaved Caspase-9 and cleaved Caspase-3 confirm activation of the mitochondrial apoptotic pathway (n = 3). Data are presented as median with 95% confidence interval. \**P* < 0.05, \*\**P* < 0.01, \*\*\**P* < 0.001; ns, not significant. Data in (E) to (G) were analyzed using the Mann-Whitney U test.

Transmission electron microscopy (TEM) subsequently revealed classic apoptotic bodies and extensive mitochondrial damage in testicular stromal cells, including mitochondrial swelling, cristae disruption, and vacuolization (Fig. 5D). In contrast, no obvious mitochondrial abnormalities and apoptotic bodies were observed in hepatocytes (Fig.S4F). To investigate the underlying mechanism, we isolated primary testicular stromal cells and performed Western blotting analysis. This analysis showed that *V. dispar* treatment decreased mitochondrial cytochrome C (CytC), increased expression of the pro-apoptotic protein Bax, reduced anti-apoptotic Bcl-2, and elevated levels of cleaved Caspase-9 and cleaved Caspase-3 (Fig. 5E–G). These findings collectively indicate the activation of the intrinsic mitochondrial apoptosis pathway.

### 3.6 *V. dispar* gavage and sequential administration of liver-specific antigen S100 effectively simulated typical AIH

The experimental results described above indicate that *V. dispar* induces significant proliferation and infiltration of CD19^+^ B cells in the liver. To investigate the potential role of *V. dispar*-induced proliferation and infiltration of B cells in the progression of AIH and PBC, female clean-grade and germ free mice aged 6–8 weeks were treated with *V. dispar* or PBS via oral gavage. All experimental procedures followed those previously described. Following 8 weeks of gavage with either PBS or *V. dispar*, a low dose of S100 (0.25 mg) was administered intraperitoneally (Fig. 6C and Fig. S9C). Week 8 after *V. dispar* or PBS gavage was designated as day 0. Serum levels of ALT, AST, antinuclear antibodies (ANA), and IgG were measured on day 14 and day 28. Mice treated with *V. dispar* gavage for 12 weeks and low-dose S100 immunization for 28 days showed significantly higher serum ALT and AST levels in the *V. dispar* + S100 group compared to the PBS + S100 group (Fig. 6A and Fig. S9A). Compared with the PBS + S100 group, the *V. dispar* + S100 group exhibited significantly elevated serum IgG levels and a notable upward trend in ANA levels (Fig. 6B and Fig. S9B). Furthermore, H&E staining of liver tissues from both clean-grade and germ-free mice revealed that the combination of *V. dispar* gavage and S100 immunization exacerbated portal lymphocyte infiltration and interface hepatitis (Fig. 6D-F). Flow cytometric results revealed that *V. dispar* gavage markedly increased the proportion of intrahepatic B cells. Although an increased B cell proportion was also observed in the PBS + S100 group, the *V. dispar* + S100 group exhibited an even greater increase (Fig. 6G and Fig. S9D). Following S100 immunization, intrahepatic CD3^+^ T cells and CD3^+^CD4^+^ T cells proliferated markedly, with proliferation being most pronounced in the *V. dispar* + S100 group (Fig. 6H, I and Fig. S9 E, F).

**Figure 6.**
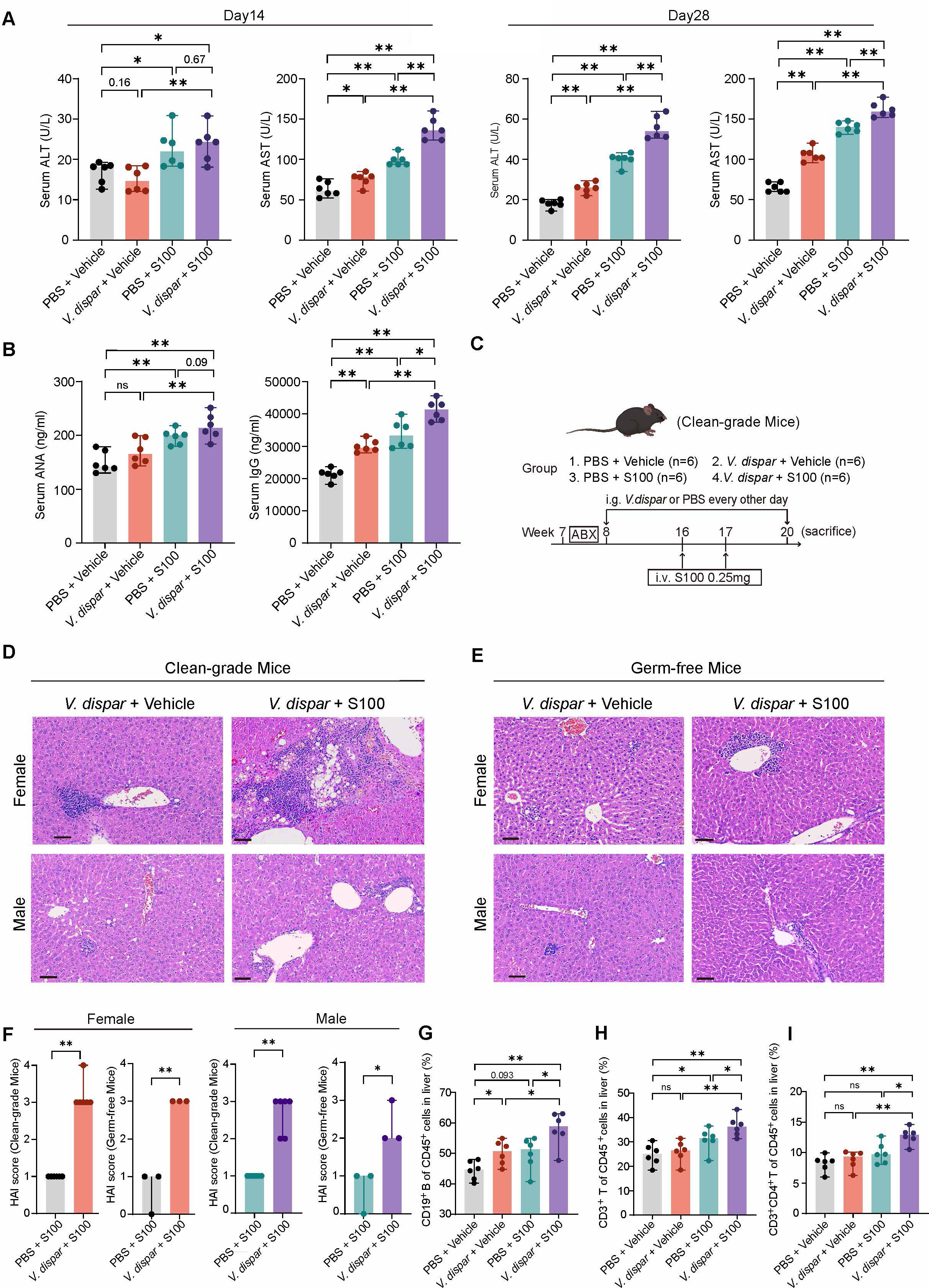
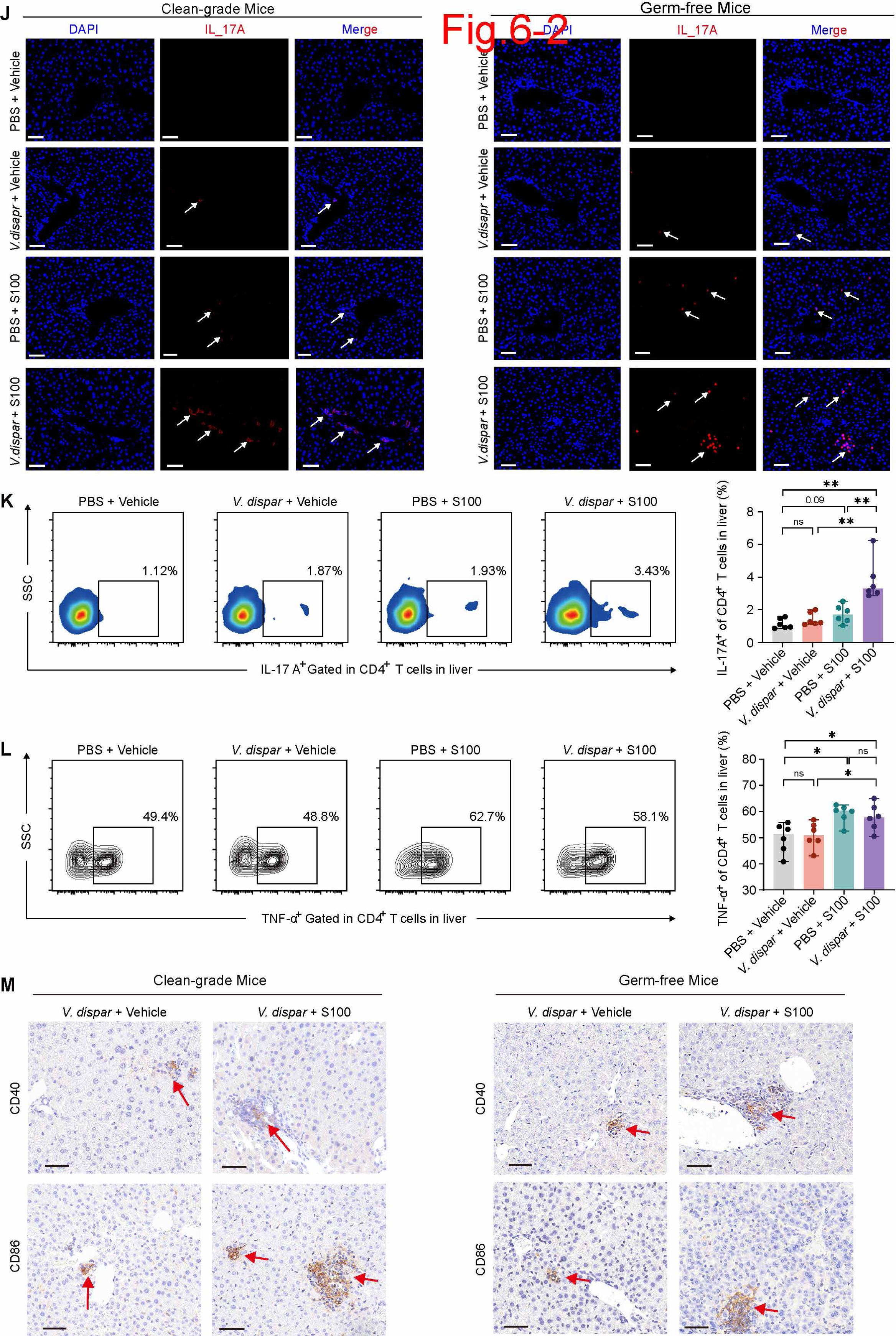
*V. dispar* plus hepatic-specific antigen S100 administration elicits an AIH-like phenotype and promotes Th17-cell activation. (A) Serum ALT and AST levels determined on days 14 and 28 post-S100 injection (n = 6). (B) Serum autoantibodies ANA and IgG measured by ELISA on day 28 post-S100 injection (n = 6). (C) Experimental design: mice were gavaged with *V. dispar* or PBS for 8 weeks, then received low-dose S100 intraperitoneal injection or adjuvant alone, with continued gavage until week 20. (D, E) Representative H&E-stained liver sections from male and female clean-grade (D) and GF (E) mice. Scale bar: 60 μm. (F) Liver injury scored by HAI (n = 6). (G–I) Analysis of liver-infiltrating CD19^+^ B cells (G), CD3^+^ T cells (H), and CD3^+^CD4^+^ T cells (I) (n = 6). (J) Representative immunofluorescence images of DAPI and IL-17A in the liver. Scale bar: 50 μm. (K, L) Representative flow cytometric plots and analysis of IL-17A^+^ (Th17) cells (K) and TNF-α^+^ cells (L) among liver-infiltrating CD4^+^ T cells (n = 6). (M) Representative immunohistochemical staining images of CD40 and CD86 proteins in the liver. Data are presented as median with 95% confidence interval. \**P* < 0.05, \*\**P* < 0.01, \*\*\**P* < 0.001; ns, not significant. Data in (A, B, G–I, K, and L) were analyzed using the Kruskal-Wallis test with Dunn’s multiple comparisons correction; for (F) using the Student’s t-test.

Th17 cells play an important role in AIH and PBC. Immunofluorescence staining of livers from clean-grade and germ-free mice revealed significantly elevated hepatic IL-17 cytokine levels after 12 weeks of *V. dispar* gavage and 28 days of low-dose S100 immunization, compared with the *V. dispar* alone or low-dose S100 alone groups (Fig. 6J). Flow cytometry demonstrated a significant increase in the proportion of IL-17A^+^ cells (Th17 cells) among hepatic CD4^+^ T cells (Fig. 6K). In the PBS+S100 and *V.dispar* +S100 groups, the proportion of TNF-α^+^ cells among CD4^+^ T cells markedly increased compared to the PBS+Vehicle and *V. dispar* + Vehicle groups; however, the proportion of TNF-α^+^ cells did not significantly differ between the PBS + S100 and *V. dispar* + S100 groups (Fig. 6L). Immunohistochemical analysis demonstrated significantly elevated levels of the costimulatory molecules CD40 and CD86 in the livers of both clean-grade and germ-free mice from the *V. dispar*+S100 group (Fig. 6M). These results indicate that *V. dispar* gavage induces the proliferation of B cells in the liver while S100 upregulates the levels of CD40 and CD86. Stimulated by the liver-specific antigen S100, Th17 cell activation was promoted. The gating strategy for flow cytometry is shown in Fig. S9H and S9I.

### 3.7 *V. dispar* gavage and sequential administration of PDC-E2 mimotope effectively simulated typical PBC

Female clean-grage and germ-free C57BL/6J mice aged 6–8 weeks were administered *V. dispar* or PBS via gavage, followed by intraperitoneal injection of a PDC-E2 mimotope (Fig. 7D and Fig. S10A). Serum biochemical parameters and levels of AMA-M2 and IgM autoantibodies were measured on day 21 and day 42. Our results showed that after 14 weeks of *V. dispar* gavage and 42 days of peptide immunization, serum levels of ALP, TBA, and TC in the *V. dispar* + peptide group were significantly higher than those in the *V.dispar* + Vehicle group (Fig. 7A, B). Compared with the *V.dispar* + Vehicle group, the *V. dispar* + peptide group showed significantly elevated serum levels of AMA-M2 and IgM autoantibodies (Fig. 7C, and Fig. S10B). Furthermore, H&E staining of liver tissues from both clean-grade and germ-free mice revealed that *V. dispar* gavage combined with peptide immunization exacerbated portal lymphocyte infiltration and bile duct injury (Fig. 7E, F). Flow cytometry results showed that peptide treatment significantly increased the proportion of hepatic CD3^+^ T cells, whereas the proportion of B cells showed no obvious change. Following peptide immunization, intrahepatic CD3^+^ T cells proliferated markedly, with an even more pronounced increase in the *V. dispar* + peptide group (Fig. 7G, H, and Fig. S10C, D). Immunofluorescence staining of liver tissues from clean-grade and germ-free mice demonstrated that *V. dispar* gavage upregulated hepatic IL-17 levels in the peptide-treated group (Fig. 7I). Flow cytometric analysis demonstrated a significant increase in the proportion of hepatic Th17 cells among CD4^+^ T cells following peptide treatment, an effect further augmented by *V. dispar* gavage (Fig. 7J). Immunohistochemical analysis revealed significantly elevated levels of the costimulatory molecules CD40 and CD86 in the livers of both clean-grade and germ-free mice from the *V. dispar* + peptide group (Fig. 7K). In summary, these results further elucidated that *V. dispar* gavage-induced proliferation of intrahepatic B cells while PDC-E2 mimotope importantly contributes to Th17 cell activation by providing essential auxiliary support.

**Figure 7.**
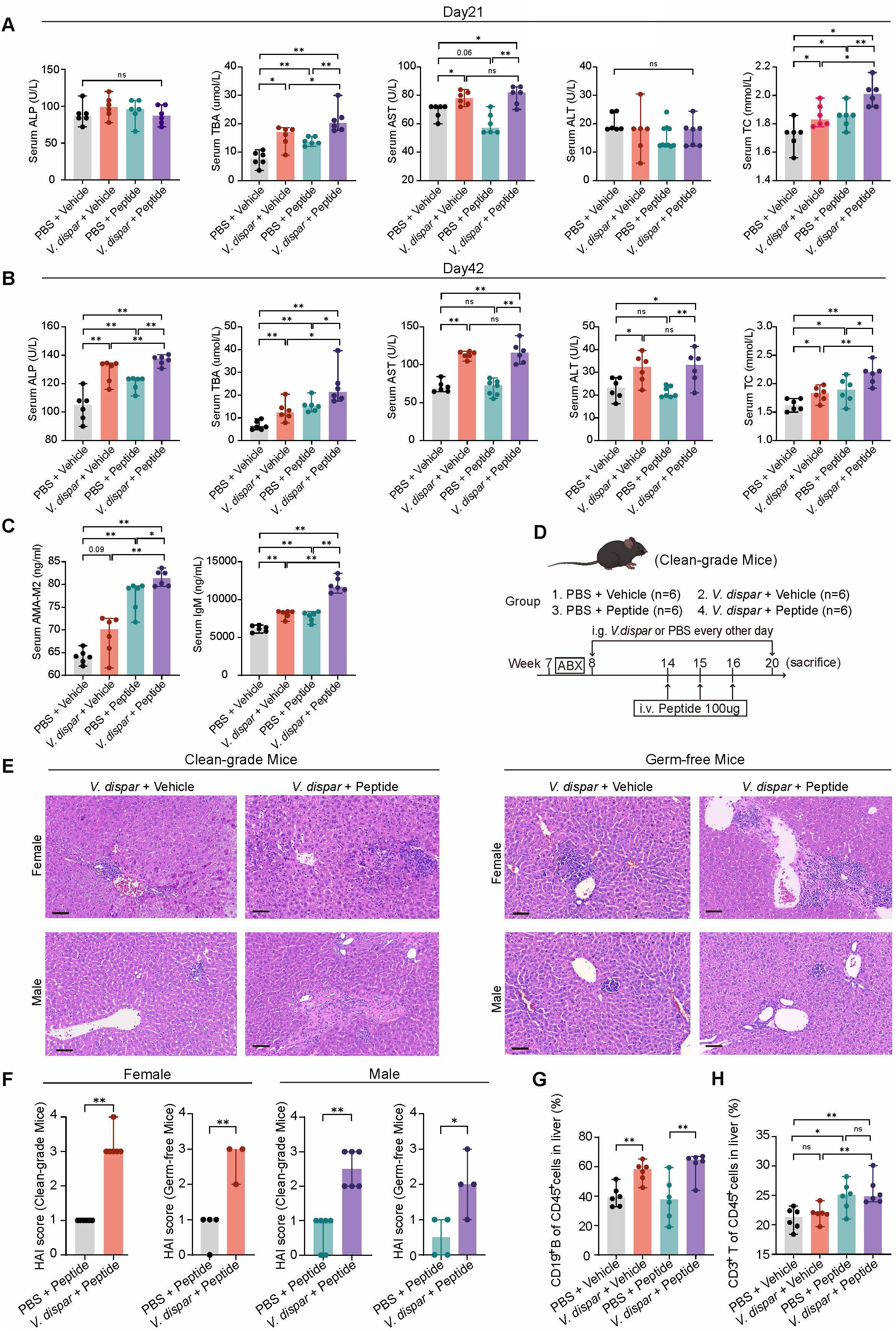
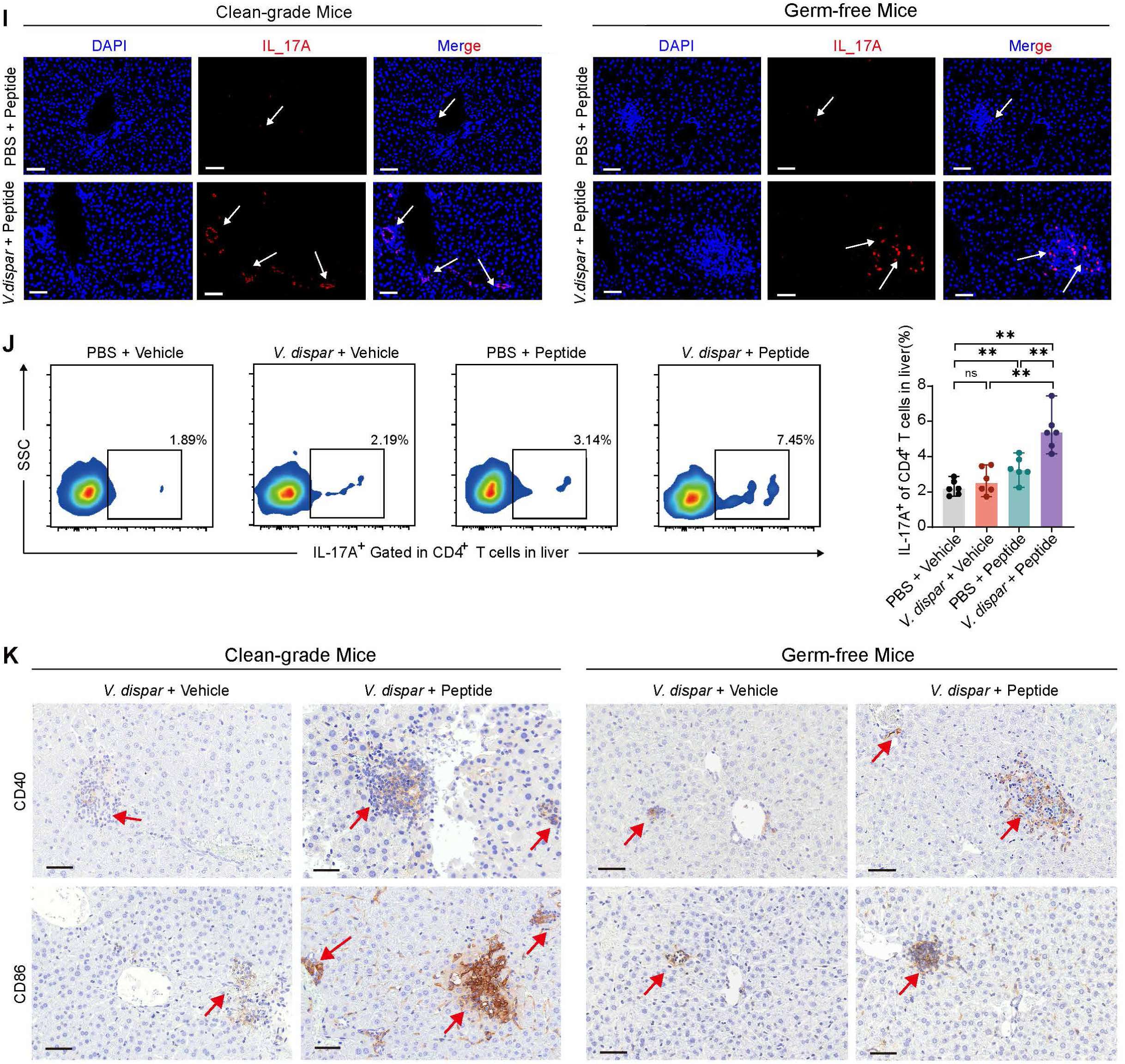
*V. dispar* plus PDC-E2 mimetic peptide administration elicits a PBC-like phenotype and promotes Th17-cell activation. (A, B) Serum biochemistry detected on days 21 (A) and 42 (B) post-PDC-E2 injection (n = 6). (C) Serum AMA and IgM levels measured by ELISA on day 42 post-PDC-E2 injection (n = 6). (D) Experimental design: Mice were gavaged with *V. dispar* or PBS for 6 weeks, then received PDC-E2 mimetic peptide or adjuvant alone. Gavage was continued until week 20. (E) Representative H&E stained liver sections from male and female clean-grade and GF mice. Scale bar: 60 μm. (F) HAI based liver injury scoring (n = 3–6). (G, H) Quantification of liver-infiltrating CD19^+^B cells (G) and CD3^+^ T cells (H) (n = 6). (I) Representative immunofluorescence images of DAPI and IL-17A in the liver. Scale bar: 50 μm. (J) Representative flow cytometric plots and analysis of IL-17A^+^ (Th17) cells among liver-infiltrating CD4^+^ T cells (n = 6). (K) Representative immunohistochemical staining images of CD40 and CD86 in livers from clean-grade and GF mice. Scale bar: 50 μm. Data are presented as median with 95% confidence interval. \**P* < 0.05, \*\**P* < 0.01, \*\*\**P* < 0.001; ns, not significant. Data in (A–C, G, and H) were compared using the Kruskal-Wallis test with Dunn’s multiple comparisons correction. For (F), the Student’s t-test was used.

### 3.8 Altered *V. dispar*-testosterone-BAFF axis and serum biomarkers in treatment-naïve AIH and PBC patients

Multiplex flow immunofluorescence assays demonstrated significantly elevated serum IL-17 in both patient groups, indicative of Th17 cell polarization (Fig. 8A). Marked increases in a broad spectrum of cytokines further pointed to concomitant Th1 (Fig. 8B–E) and Th2 (Fig. 8F–H) cell activation, alongside a heightened systemic inflammatory milieu (Fig. 8I–L). Of note, the relative abundance of *V. dispar* in fecal samples correlated inversely with serum L-arginine, testosterone, while positively correlated with BAFF levels (Fig. 8M–O). Moreover, serum arginine and testosterone were significantly lower in PBC and AIH patients relative to healthy controls, whereas BAFF was significantly higher (Fig. 8P–R).

**Figure 8.**
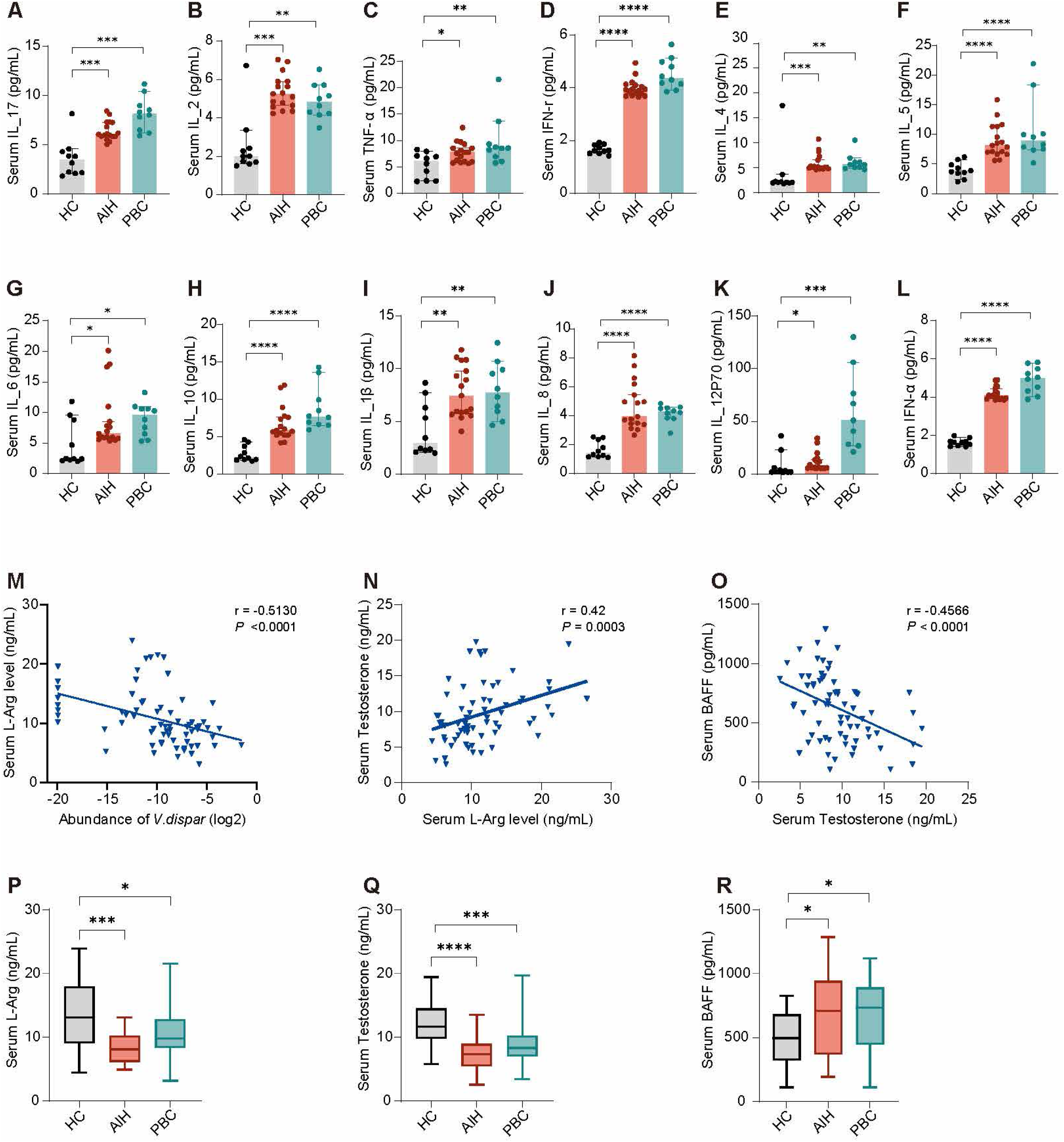
Cytokine profiles and correlation analyses in treatment-naïve AIH and PBC patients. (A–L) Serum cytokine levels (n = 37). (M) Fecal *V. dispar* abundance was negatively correlated with serum L–arginine (n = 70). (N) Serum L–arginine was positively correlated with serum testosterone (n = 70). (O) Serum testosterone was negatively correlated with serum BAFF (n = 70). (P–R) Serum arginine and testosterone levels were decreased in both the AIH and PBC groups, whereas serum BAFF levels were increased. Data are presented as median with 95% confidence interval (n = 70). \**P* < 0.05, \*\**P* < 0.01, \*\*\**P* < 0.001; ns, not significant. Data in (A–L, and P–R) were compared using the Kruskal-Wallis test with Dunn’s multiple comparisons correction. Statistical analyses for (M–O) were conducted using Spearman’s rank correlation.

## 4. Discussion

In summary, we found that the initiation and development of AIH and PBC unfold in two distinct stages. In the first stage, systematic hypoargininemia, induced by *V. dispar*, led to mitochondrial apoptosis of gonadal stromal cells. The significantly lowered testosterone levels in mice, which in turn weakened testosterone’s inhibitory effect on BAFF secretion. Consequently, B-cell infiltration around the portal tracts and bile duct was aggravated, leading to the development of AIH/PBC-like pathological manifestations. In the second stage, the induction of the liver-specific antigen S100 and the PDC-E2 homologous mimotope facilitated the activation of B cells and Th17 cells. This further resulted in the typical interface hepatitis and bile duct injury characteristic of AIH and PBC, respectively.

Our findings also shed light on the prominent clinical feature of AIH and PBC: their predilection for females, the mechanism of which has been unclear. We propose that the low testosterone levels observed in the first stage, stemming from gonadal stromal cell destruction caused by *V. dispar*, play a critical role. Research has shown that AIH exhibits some characteristic changes in gut bacteria and metabolite profiles. It is reported that *Veillonella* is significantly enriched whereas arginine is significantly underrepresented in the AIH patients from Egypt and China [19, 27]. Similarly, the gut microbiome from the PBC patients is characterised by under-representation of proteins involved in arginine and proline metabolism [17]. Consistent with these findings, we also found that arginine levels were significantly reduced in the *V. dispar* treated group including both clean-grade and germ-free mice. Therefore, it seems reasonable to infer that specific bacteria and related metabolites may be involved in the pathogenesis of AIH.

Our study indicated that low testosterone level caused by intestinal *V. dispar* aggravated intrahepatic B-cell infiltration, contributing to AIH/PBC-like pathologies. Thus, testosterone appears to have a protective effect against the development of AIH and PBC. Further supporting this, Fig. 4C demonstrates that female mice consistently exhibit lower serum testosterone levels than male mice, regardless of *V. dispar* treatment. This hormonal difference provides a partial explanation for the observed female preference in the development of AIH and PBC. Our results align with previous reports on the protective effects of testosterone in animal models of multiple sclerosis and SLE [28, 29]. Testosterone treatment ameliorated hepatic inflammation in experimental cholangitis [30]. Furthermore, clinical studies have demonstrated that testosterone therapy ameliorates symptoms in women with autoimmune diseases, including SLE, rheumatoid arthritis, multiple sclerosis, and Sjögren’s syndrome [31–33]. Sex-based differences in the prevalence of AIH and PBC constitute their most prominent clinical feature, as both conditions disproportionately affect females. However, the underlying mechanism remains unclear. This study found that low testosterone level induced by intestinal *V. dispar* contributed to the development and female preference of AIH and PBC.

Traditionally, both AIH and PBC are regarded as autoimmune diseases primarily driven by T cells. However, evidence also supports a critical role for B cells in the pathogenesis of AIH and PBC. Patients with AIH typically exhibit higher levels of IgG, while those with PBC often show elevated IgM titers. Both IgG and IgM are produced by plasma cells, which are mature B cells. Furthermore, previous studies have revealed significant B cell infiltration in the livers of both PBC and AIH patients [34–37]. Effects of rituximab, a B-cell-depleting therapy, on AIH also suggest that B cells play a critical role in the induction and progression of the disease [38]. In our study, BAFF-neutralization blocked the *V. dispar*-mediated exacerbation of AIH or PBC-like pathology, further indicating the role of B cells in the pathogenesis of AIH and PBC.

In the second stage, B cells and Th17 cells were activated by the liver-specific antigen S100 and the PDC-E2 homologous mimotope. In our study, we found that *V. dispar* + S100 or *V. dispar* + PDC-E2 mimotope together upregulated the expression of co-stimulatory molecules CD40 and CD86, indicating the role of S100 or PDC-E2 mimotope in activating B cells and T cells. Furthermore, our experimental results demonstrated that while oral administration of *V. dispar* alone induced predominant portal B cell infiltration, T cell activation and typical histopathological features of AIH or PBC were not observed. Further, we found that oral administration of *V. dispar* alone induced predominant portal B cell activation, but no T cell activation or typical histopathological features of AIH or PBC. In striking contrast, with the aid of S100 or PDC-E2 mimetic peptide, oral administration of *V. dispar* induced elevated serum levels of autoantibodies (ANA and AMA-M2) and significantly increased proportions of hepatic IL-17A^+^ CD4 ^+^ T cells. Th17 cells, characterized by the secretion of IL-17, accumulate around bile ducts undergoing chronic nonsuppurative destructive cholangitis, indicating their role in the pathogenesis of PBC [39]. PDC-E2 is widely recognized as the primary autoantigen target in PBC [40]. The liver-specific antigen S100 has been implicated in the induction of experimental autoimmune hepatitis probably mediated by autoreactive T cells [24]. In the presence of S100 or PDC-E2 mimotope, *V. dispar* treated mice exhibited canonical pathological hallmarks of AIH and PBC, respectively.

Overall, the pathogenesis of AIH and PBC involves a clear “dual-hits” process. Non-specific B cell proliferation and intrahepatic infiltration induced by *V. dispar* or other factors represent the first hit, while the second hit entails the activation of B cells and T cells triggered by autoreactive antigens such as S100 or PDC-E2. The first hit clearly delineates numerous similarities between AIH and PBC, including female predominance, significantly elevated intestinal *V. dispar* levels, increased serum IgG or IgM, and intrahepatic B cell infiltration. Furthermore, AIH and PBC frequently overlap in clinical manifestations. The second hit explains why AIH and PBC, despite a shared pathogenic basis, exhibit distinct clinical manifestations and hepatic pathological features. For instance, AIH mainly presents with elevated ALT, and AST, whereas PBC typically shows elevated GGT, and ALP. Histopathologically, AIH primarily manifests as interface and lobular hepatitis; while PBC mainly presents as florid bile duct lesions and ductopenia.

By examining the interaction between a specific bacterial infection and the host immune system, our study provides a novel paradigm for a deeper understanding of the pathogenesis of AIH and PBC, which can be extended to other infection-associated autoimmune diseases. However, this study has several limitations. First, although BAFF was targeted with a neutralizing antibody, the direct role of testosterone was not assessed in BAFF-overexpressing or knockout models. Second, the complexity of metabolic networks within the gut microbiota suggests that metabolites beyond arginine may also mediate testosterone reduction. Third, the proposed *V. dispar*–arginine–testosterone–BAFF axis requires validation in large-scale clinical cohorts to confirm its translational potential.

## Supporting information

Supplementary Methods, Results, Figures

## SUPPLEMENTAL INFORMATION

Supplemental information can be found online at

## ACKNOWLEDGMENTS

We sincerely thank Prof. Xiong Ma of Division of Gastroenterology and Hepatology, Renji Hospital, School of Medicine, Shanghai Jiao Tong University, for kindly providing some of the clinical samples. This study was supported by the Natural Science Foundation of China (no. 82372318), and Major Research Project for Young and Middle-aged People of the Health Commission of Fujian Province (no. 2022ZQNZD004).

## AUTHOR CONTRIBUTIONS

Y.L.L., Q.Q.X., and X.L. performed the experiments and drafted the manuscript. W.W.Z, Y.H.C., Y.Y.Y., L.N.Z., X.N.C., M.X.W., and H.X.X. assisted with the mouse experiments. J.S.P. provided critical comments and contributed to manuscript revision. S.W., and Y.L.L. also designed and supervised the study and finalized the manuscript.

---

**Figure S1. *V. dispar* increases splenic B cells without altering intrahepatic CD3^+^ T cells**

(A) C57BL/6 mice were gavaged with *V. dispar* or PBS for 12 weeks. (B) Hepatic CD3^+^ T cell frequency (CD3^+^/CD45^+^) showed no significant intergroup differences in either sex (n = 6). (C) Representative flow cytometric plots showing significantly elevated splenic CD19^+^ B cells (CD19^+^/CD45^+^) in *V. dispar*-treated mice of both sexes (n = 6). (D, E) No significant differences were observed in splenic CD3^+^ T cell and its subset frequencies in either sex (n = 6). (F) Representative immunofluorescence images of CD19 (green) indicating B cell infiltration in the liver and spleen of SPF mice. Data are presented as median with 95% confidence interval. \**P* < 0.05, \*\**P* < 0.01, \*\*\**P* < 0.001; ns, not significant. Data in (B) to (E) were analyzed using the Mann-Whitney U test.

**Figure S2. BAFF mediates *V. dispar*-induced B cell expansion and liver injury**

(A) Representative immunofluorescence showing increased BAFF (red) in the *V. dispar* group and its reduction by anti-BAFF. Scale bar: 100 μm. (B) Representative immunohistochemistry showing increased hepatic CD19^+^ B cells in *V. dispar*-treated mice and their reduction by anti-BAFF treatment. Scale bar: 100 μm. (C) Representative flow cytometric plots showing increased splenic CD19^+^ B cells in *V. dispar*-treated mice and partial reversal with anti-BAFF (n = 6). (D) Anti-BAFF reduced serum ALP and AST levels in both sexes (n = 6). Data are presented as median with 95% confidence interval. \**P* < 0.05, \*\**P* < 0.01, \*\*\**P* < 0.001; ns, not significant. Data in (C) and (D) were analyzed using the Kruskal-Wallis test.

**Figure S3. Testosterone supplementation reverses *V. dispar*-induced BAFF elevation and B cell infiltration in non-immunized male mice**

(A, B) Serum testosterone levels were reduced in ORX + *V. dispar* mice and restored by testosterone supplementation, with data shown for males (A) and females (B) (n = 6). (C, D) Representative flow cytometric plots showing increased splenic CD19^+^ B cells in ORX + *V. dispar* mice, with partial reduction by testosterone in males (C) and females (D) (n = 6). Data are presented as median with 95% confidence interval. \**P* < 0.05, \*\**P* < 0.01, \*\*\**P* < 0.001; ns, not significant. Data in (A) and (D) were analyzed using the Kruskal-Wallis test.

**Figure S4. *V. dispar* disrupts gonadal architecture and induces ultrastructural damage in testicular stromal cells**

(A) Testes-to-body weight ratios showed no significant intergroup differences. (B) Representative H&E staining of testes showing reduced germ cells and disorganized interstitium in *V. dispar*-treated mice (n = 6). Scale bar: 100 μm. (C) Representative immunofluorescence of cleaved Caspase-3 (green) in ovaries from GF mice showing increased apoptosis after *V. dispar* treatment. DAPI (blue) labels nuclei. Scale bar: 100 μm. (D, E) Representative immunohistochemistry and immunofluorescence images of 3β-HSD showing diminished 3β-HSD signals in Leydig/stromal cells of *V. dispar*-treated groups. Scale bar: 100 μm. (F) Transmission electron microscopy analysis of mitochondrial ultrastructure in hepatocytes. Data are presented as median with 95% confidence interval. \**P* < 0.05, \*\**P* < 0.01, \*\*\**P* < 0.001; ns, not significant. Data in A were analyzed using the Mann-Whitney U test.

**Figure S5. *V. dispar*-expressed arginine decarboxylase (ADC) promotes arginine catabolism and exacerbates AIH/PBC-like pathogenesis**

(A, B) Serum metabolomics revealed distinct metabolic profiles between PBS and *V. dispar* groups, characterized by significant enrichment of the arginine (Arg) metabolism pathway (A) and reduced serum arginine levels (B) (n = 6). (C) Representative H&E staining of testes showing improved interstitial structure after arginine supplementation in *V. dispar*-treated mice. Scale bar: 100 μm. (D) Representative immunofluorescence staining of testes indicating a preserved stromal structure and suppressed apoptosis after arginine supplementation in *V. dispar* treated mice. Scale bar: 100 μm. (E, F) Arginine supplementation restored serum testosterone and reduced serum BAFF levels (n = 6). (G, H) Representative flow cytometric plots showing reduced frequencies of intrahepatic and splenic CD19^+^B cells after arginine supplementation (n = 6). (I-L) Arginine supplementation lowered serum AST, and ALP without affecting TC or ALT (n = 6). Data are presented as median with 95% confidence interval. \**P* < 0.05, \*\**P* < 0.01, \*\*\**P* < 0.001; ns, not significant. Data in (B) were analyzed using the Mann-Whitney U test; for (E) to (L) using the Kruskal-Wallis test with Dunn’s multiple comparisons correction.

**Figure S6. Arginine supplementation or ADC-expressing *E. coli* modulates AIH/PBC-like pathogenesis**

(A, B) NCBI genomic analysis identified the *speA* gene encoding ADC in *V. dispar*. A recombinant ADC-expressing *E. coli* strain (pET28a-*speA*/BL21) was performed and validated by agarose gel electrophoresis. (C) C57BL/6 mice were gavaged with control *E. coli* or recombinant *E. coli-speA*. (D-F) Serum Arg and testosterone levels were significantly decreased in the *E. coli-speA* group (n = 6). (G) Representative H&E staining of testes showing aggravated interstitial damage in *E. coli*-*speA* treated mice. Scale bar: 100 μm. (H, I) *E. coli*-*speA* elevated serum and hepatic BAFF levels (n = 6). (J, K) *E. coli*-*speA* gavage increased CD19^+^ B cells in the liver (J) and spleen (K), recapitulating the phenotype induced by *V. dispar* (n = 6). Data are presented as median with 95% confidence interval. \**P* < 0.05, \*\**P* < 0.01, \*\*\**P* < 0.001; ns, not significant. \**P* < 0.05, \*\**P* < 0.01, \*\*\**P* < 0.001; ns, not significant. Data in (D) to (E) and (H) to (K) were analyzed using the Mann-Whitney U test.

**Figure S7. Arginine deficiency impairs mitochondrial function and suppresses testosterone biosynthesis in testicular stromal cells**

(A) GO enrichment analysis showing “negative regulation of apoptotic process” as a downregulated term. (B) Counts of up/downregulated DEGs in pathway. (C) GSEA plot of the negative regulation of apoptotic process pathway. (D) Transmission electron microscopy of TM3 cells showing mitochondrial swelling, cristae loss, and vacuolization after 48 h of arginine deprivation. Scale bar: 5.0 μm (left), 500 nm (right). (E) Apoptosis (Annexin V^+^/PI^+^) induced by arginine deficiency was attenuated by the caspase inhibitor Z-VAD-FMK, but not by Nec-1 or Liproxstatin-1 (n = 3). (F) Flow cytometric detection of intracellular ROS accumulation after 24 or 48 h of arginine deprivation (n = 3). (G) JC-1 staining showing loss of mitochondrial membrane potential (ΔΨm) in arginine-deprived cells (n = 3). (H, I) Representative immunofluorescence showing time-dependent CytC release from mitochondria to cytosol after arginine deprivation. The cells were stained with mitochondrial probes (Mito-Tracker Red CMXRos), incubated with anti-CytC (Green) and then observed by laser confocal microscopy. Pearson’s correlation coefficient was used to assess the colocalization between mitochondria and cytochrome c. The decreased Pearson’s correlation coefficient upon arginine deprivation reflects cytochrome c (CytC) release from mitochondria into the cytosol, which increases over time (n = 3). Scale bar: 10 μm. (J) Western blotting showing reduced steroidogenic enzymes P450scc and StAR after 48 h of arginine deprivation (n = 3). Data are presented as median with 95% confidence interval. \**P* < 0.05, \*\**P* < 0.01, \*\*\**P* < 0.001; ns, not significant. Data in (E) to (I) were performed using the Kruskal-Wallis test; for (J) using the Mann-Whitney U test.

**Figure S8. Arginine deficiency disrupts oxidative phosphorylation and reduces histone H3 acetylation**

(A) KEGG pathway enrichment analysis revealing oxidative phosphorylation as the top dysregulated pathway under arginine deprivation. (B) qRT-PCR showing suppression of oxidative phosphorylation-related genes (COX5A, ATP5D, UCP2, HSD17B4, NDUFS3, UQCRB) in arginine-deprived TM3 cells (n = 3). (C) Enzymatic assays showing reduced activities of mitochondrial complexes I, II, IV, and V in the absence of arginine (n = 3). (D) Western blotting showing reduced multi-site acetylation of histone H3 (K4 + K9 + K14 + K18 + K23 + K27) under arginine deprivation (n = 3). Data are presented as median with 95% confidence interval. \**P* < 0.05, \*\**P* < 0.01, \*\*\**P* < 0.001; ns, not significant. Data in (B) to (D) were analyzed using the Mann-Whitney U test.

**Figure S9. *V. dispar* plus low-dose S100 immunization exacerbates portal lymphocyte infiltration and interface hepatitis**

(A) Serum ALP, TC, and TBA levels measured on days 14 and 28 post-S100 injection (n = 6). (B) Serum ANA levels in GF mice quantified using ELISA on day 28 post-S100 injection (n = 6). (C) Experimental design in GF mice. (D–G) Representative flow cytometric plots of liver-infiltrating cells: CD19^+^B cells (D), CD3^+^T cells (E), CD3^+^CD4^+^T cells (F), and CD3^+^CD8^+^T cells (G). (H) Gating strategy for hepatic CD19^+^, CD3^+^, CD3^+^CD4^+^, and CD3^+^CD8^+^ cells (CD45^+^-gated). (I) Gating strategy for hepatic TNF-α^+^ cells. Data are presented as median with 95% confidence interval. \**P* < 0.05, \*\**P* < 0.01, \*\*\**P* < 0.001; ns, not significant. Data in (A) and (B) were analyzed using the Kruskal-Wallis test.

**Figure S10. *V. dispar* plus PDC-E2 mimotope immunization exacerbates PBC-like phenotype and Th17 activation**

(A) Experimental design in GF mice. (B) Serum AMA-M2 levels in GF mice detected via ELISA on day 42 post-PDC-E2 injection (n = 3). (C, D) Representative flow cytometric plots of liver-infiltrating CD19^+^ B cells and CD3^+^ T cells. Data are presented as median with 95% confidence interval. \**P* < 0.05, \*\**P* < 0.01, \*\*\**P* < 0.001; ns, not significant. Data in (B) were analyzed using the Kruskal-Wallis test.

