## Supplementary Methods, Results, Figures for "*Veillonella dispar* triggers “dual-hits” in the pathogenesis of autoimmune hepatitis and primary biliary cholangitis via testosterone depletion-BAFF-B cell axis and Th17 cell activation"

### *2. Materials and methods*

#### 2.1 Extraction of S100 Liver Antigen

Livers were perfused with pre-cooled phosphate-buffered saline until they appeared pale, and then homogenized in pre-chilled phosphate-buffered saline. After repeated freeze-thaw cycles at -80 °C, the liver homogenates were centrifuged at 150×g for 15 minutes at 4 °C. The supernatant was collected and subsequently centrifuged at 100,000×g for 1 hour in an ultracentrifuge. The supernatant containing S100 was harvested and concentrated using an Amicon® Ultra-15 centrifugal filter unit (Millipore, Burlington, MA, USA). Separation was then performed using an AKTA Pure system (GE Healthcare, Chicago, IL, USA) equipped with a 90-cm CL-6B Sepharose® chromatography column (Pharmacia & Upjohn Company LLC, Kalamazoo, MI, USA). The first non-toxic elution peak was collected, and its concentration was adjusted to 0.5g-2.0g/L.

#### 2.2 Culture of *V.dispar*

*V.dispar* (ATCC 17748) was cultured in Veillonella Selective Medium (VS medium) following to the American Type Culture Collection (ATCC, American) guidelines. Bacteria were cultured under anaerobic conditions at 37°C for 24 hours. *V. dispar* was confirmed via 16S rDNA sequencing (V4 sequences). The complete genome sequence was performed by Tsingke (Beijing, China).

#### 2.3 Serologic assays

Serum levels of alkaline phosphatase (ALP), alanine aminotransferase (ALT), AST, total cholesterol (TC), and triglyceride (TG) were determined using an automatic biochemistry analyzer (Cobas8000, Roche Diagnostics, Switzerland) according to the manufacturer's instructions. Serum immunoglobulin M (IgM) and BAFF were quantified using ELISA kits (Beyotime, PI489 and PB062, respectively; Shanghai, China). The serum arginine levels were determined using an ELISA kit (MLBIO, 13533, Shanghai, China).

#### 2.4 Real-time quantitative PCR

Total RNA was extracted using TRIzol reagent (Invitrogen, 15596-026, CA, USA) and reverse-transcribed into cDNA with a reverse transcription kit (Yeasen, 11141ES60, Shanghai, China). Relative mRNA expression levels of the target genes were subsequently quantified using SYBR Green PCR Master Mix (Yeasen, 11201ES08, Shanghai, China) on a QuantStudio5 Real-time quantitative PCR detection system (Applied Biosystems, USA), with normalization to β-actin. Primer sequences, synthesized by Tsingke (Beijing, China), are provided in Table S1.

#### 2.5 Western blotting

Total protein was extracted following lysis of tissues or cells in a lysis buffer supplemented with a cocktail of protease inhibitors (Epizyme, GRF101, Shanghai, China) and phosphatase inhibitors (Epizyme, GRF102, Shanghai, China). Protein concentrations were quantified using a bicinchoninic-acid (BCA) protein assay kit

(Epizyme, ZJ102, Shanghai, China). Protein samples were mixed with SDS-PAGE loading buffer (Epizyme, LT101, Shanghai, China), separated by 8%–10% SDS-PAGE, and subsequently transferred to polyvinylidene difluoride (PVDF) membranes (Millipore, IPVH00010, MA, USA). Membranes were incubated with the primary antibody (listed in Table S2) overnight at 4 °C, followed by incubation with an HRP-conjugated secondary antibody for 1 hour at room temperature. The blots were detected using ECL (Fdbio, FD8030, Hangzhou, China).

#### 2.6 RNA sequencing

Total RNA was extracted; cDNA libraries were subsequently constructed and sequenced on an Illumina HiSeqTM 2500 (Panomix Biomedical Technology Co., Ltd., Suzhou, China). Statistical significance was defined as a $P$-value < 0.05 and a fold change > 2.

#### 2.7 Heterologous expression of speA

Standard molecular cloning techniques cloned the DNA fragment encoding full-length speA into the pET-28a vector. *E. coli*BL21 cells (Yeasen Biotechnology, 11804ES80) carrying the speA construct were grown at 37 °C to an OD600 of 0.6. Protein expression was induced with 200 μM IPTG (Selleck, S6826) for 6 h at 37 °C. Control *E. coli* and speA-overexpressing *E. coli* (*E. coli*-speA) were cultured in LB medium (Sangon Biotech, A507002) at 37 °C for 24 h.

#### 2.8 Single-cell RNA sequencing

Single-cell RNA sequencing (scRNA-seq) was performed on mouse liver tissues under different treatment conditions. After enzymatic dissociation, single-cell suspensions were prepared and loaded onto a Chromium platform (10x Genomics) for encapsulation and barcoding. Library construction and sequencing were performed by BioNovoGene (Suzhou, China) on an Illumina NovaSeq 6000 platform, with a target depth of 20,000 reads per cell. Raw sequencing data were demultiplexed and converted into FASTQ files using Illumina bcl2fastq software. The Cell Ranger pipeline (10x Genomics) was used for alignment, cell barcode recognition, and UMI quantification.

Downstream analysis was conducted using the Seurat package (version 4.0.0) in R. Quality control criteria included: genes expressed in at least three cells; 500-5000 genes detected per cell; ≥ 500 UMIs per cell; and mitochondrial gene content < 25% of total expression. Dimensionality reduction and clustering were performed using the UMAP algorithm.

#### 2.9 Metabolomic analysis

For plasma samples, 5–10 volumes of pre-chilled methanol were added, followed by vortexing and shaking. The samples were then incubated at –20 °C for 1 h and centrifuged at 14,000 × *g* for 10 min. The collected supernatant was subsequently centrifuged and filtered for further analysis.Untargeted LC–MS/MS analysis was performed using a BioNovoGene (Suzhou, China) system and a Thermo UHPLC-Q Exactive system. Metabolite identification was conducted by comparing mass spectra against entries in the HMDB (http://www.hmdb.ca/) and Metlin (https://metlin.scripps.edu/) databases. KEGG-based pathway enrichment analysis elucidated the biological significance of the differential metabolites.

The mobile phases were buffer A (95% dd$H_2O$, 5% acetonitrile, and 0.1% formic acid) and buffer B (47.5% acetonitrile, 47.5% isopropanol, 5% dd$H_2O$, and 0.1% formic acid). The gradient conditions were as follows: Positive ion mode: 0–3 min, gradient to 20% buffer B; 3–4.5 min, to 35% buffer B; 4.5–5 min, to 100% buffer B; 5–6.3 min, held at 100% buffer B; 6.3–6.4 min, returned to 0% buffer B; 6.4–8 min, held at 0% buffer B. Negative ion mode: 0–1.5 min, to 5% buffer B; 1.5–2 min, to 10% buffer B; 2–4.5 min, to 30% buffer B; 4.5–5 min, to 100% buffer B; 5–6.3 min, held at 100% buffer B; 6.3–6.4 min, returned to 0% buffer B; 6.4–8 min, held at 0% buffer B. The flow rate was maintained at 0.40 mL/min.

#### 2.10 Flow cytometry analysis

Non-parenchymal liver cells and splenic lymphocytes were isolated upon completion of the experiment. The cell suspensions were adjusted to a concentration of $1 \times 10^6$ cells/mL and centrifuged at 1000 rpm for 5 min at 4 °C. The supernatant was discarded, and the pellet washed 2–3 times with phosphate-buffered saline (PBS). Cells were incubated with Fixable Viability Stain 700 (BD Biosciences, 564997) to exclude dead cells, then stained with FITC anti-mouse CD45 (BioLegend, 103108) and PE rat anti-mouse CD19 (BioLegend, 115508) in 100 μL staining buffer for 40 min at 4 °C in the dark. Cells were washed 2-3 times with 1 mL flow cytometry staining buffer (centrifuged at 1000 rpm for 5 min, 4°C per wash). The final pellet was resuspended in 350 μL staining buffer for flow cytometric analysis. Data were processed with FlowJo software.

#### 2.11 Human sample

Treatment-naïve patients with AIH or PBC, alongside healthy controls, were enrolled from the First Affiliated Hospital of Fujian Medical University and Renji Hospital, Shanghai Jiao Tong University School of Medicine. The study was approved by the Branch for Medical Research and Clinical Technology Application, Ethics Committee of the First Affiliated Hospital of Fujian Medical University (MTCA, ECFAH of FMU No.[2025]256) and the Ethics Committee of Renji Hospital, Shanghai Jiao Tong University School of Medicine (Renji Scientific Research Ethics Review No.[2013]030). Written informed consent was obtained from all participants prior to their inclusion in the study. Stool specimens were subjected to 16S rDNA sequencing, and serum biomarkers were profiled for subjects.

#### 2.12 Statistical analyses

Statistical analyses were conducted using R software (version 4.1.3). Data are expressed as median with 95% confidence interval from at least three independent experiments. The term “n” refers to the number of biologically independent samples or animals. Specific statistical details for each experiment are provided in the corresponding figure legends. The Shapiro-Wilk test was used to assess the normality of continuous variables, followed by either the unpaired Student's t-test (for normally distributed data) or the Mann-Whitney U-test (for non-normally distributed data) to compare group differences. Comparisons among multiple groups were performed using one-way ANOVA followed by Tukey’s post hoc test for normally distributed data, or the Kruskal–Wallis test followed by Dunn’s multiple comparisons test for non-normally distributed data. Correlations were evaluated using Spearman’s rank correlation analysis. Statistical significance was defined as $P < 0.05$.

### *3. Results*

#### 3.3  *V. dispar* upregulated BAFF expression

Splenic B cell infiltration, as revealed by flow cytometry, corroborated the hepatic findings (Fig. S2C). Anti-BAFF treatment also reduced serum ALP and AST levels, consequently alleviating hepatic inflammation (Fig. S2D).

#### 3.5 *V. dispar* induced intrinsic apoptosis of gonadal stromal cells

Immunohistochemical and immunofluorescence staining for 3β-HSD in the testes revealed a significant decrease in the number of stromal cells after *V. dispar* treatment in both clean-grade and germ-free mice (Fig. S4D, and S4E).

#### 3.9 Systematic hypoargininemia caused by *V. dispar* lead to mitochondrial apoptosis of gonadal stromal cells

Serum metabolomics analysis comparing mice treated with PBS and *V. dispar* identified significant enrichment of the arginine metabolism pathway (Fig. S5A). Arginine (Arg) levels were significantly reduced in the *V. dispar* group (Fig. S5B), suggesting a potential pathological role for disrupted arginine metabolism. To further investigate, male C57BL/6 mice were divided into *V. dispar* and *V. dispar* + arginine (Arg) groups, constituting a rescue experiment designed to test the role of arginine. Testicular histology, flow cytometry of the liver and spleen, and serum analyses were performed. H&E and immunofluorescence staining showed that arginine supplementation prevented the loss of stromal cell numbers and architectural disarrangement in the testicular interstitium (Fig. S5C, and S5D). Furthermore, in the *V. dispar* + Arg group, arginine supplementation lowered serum BAFF levels, increased testosterone levels (Fig. S5E, and S5F), reduced B cell infiltration in the liver and spleen (Fig. S5G, H), and decreased serum ALP, AST, and total cholesterol levels (Fig. S5I–L).

NCBI-based genomic analysis revealed the *speA* gene in *V. dispar*, which encodes Arginine decarboxylase (ADC), an enzyme that degrades arginine. To examine its functional role, we utilized an ADC-expressing recombinant strain (pET28a-speA/BL21) (Fig. S6A, B). Male mice orally administered with *E. coli*-speA (Fig. S6C) showed significantly reduced serum arginine and testosterone levels (Fig. S6D–F). Testicular damage was also aggravated in these mice (Fig. S6G). Moreover, elevated BAFF levels were observed (Fig. S6H, I), together with increased B cell infiltration in the liver and spleen (Fig. S6J, K). These phenotypes were consistent with those observed after *V. dispar* treatment, thus supporting a role for ADC-mediated arginine catabolism in stromal dysfunction and aggravated AIH and PBC. Mitochondrial integrity is essential for testosterone synthesis in the stromal cells. Transcriptome sequencing analysis demonstrated a significant downregulation of the "negative regulation of apoptotic process" pathway in TM3 cells subjected to arginine starvation for 24 hours (Fig. S7A–C). To assess the effect of arginine, TM3 cells were cultured under arginine-sufficient or arginine-deficient conditions for 24–48 h. Arginine deprivation in TM3 cells induced apoptotic morphology including mitochondrial swelling, cristae loss, and vacuolization (Fig. S7D) and increased apoptosis, which was reversed by Z-VAD-FMK but not by Nec-1 or Liproxstatin-1 (Fig. S7E). Furthermore, arginine-deficient cells accumulated ROS (Fig. S7F), exhibited reduced $\Delta\Psi m$ (Fig. S7G), and showed time-dependent CytC release (Fig. S7H, I), all indicating intrinsic apoptosis. Additionally, arginine deprivation suppressed testosterone biosynthesis by downregulating P450scc and StAR (Fig. S7J). Transcriptomic analysis revealed altered oxidative phosphorylation pathways (Fig. S8A), with suppressed related genes (Fig. S8B), reduced complex I/II/IV/V activity (Fig. S8C), and decreased histone H3 acetylation (Fig. S8D), suggesting epigenetic involvement in these impairments.

### Fig.S1

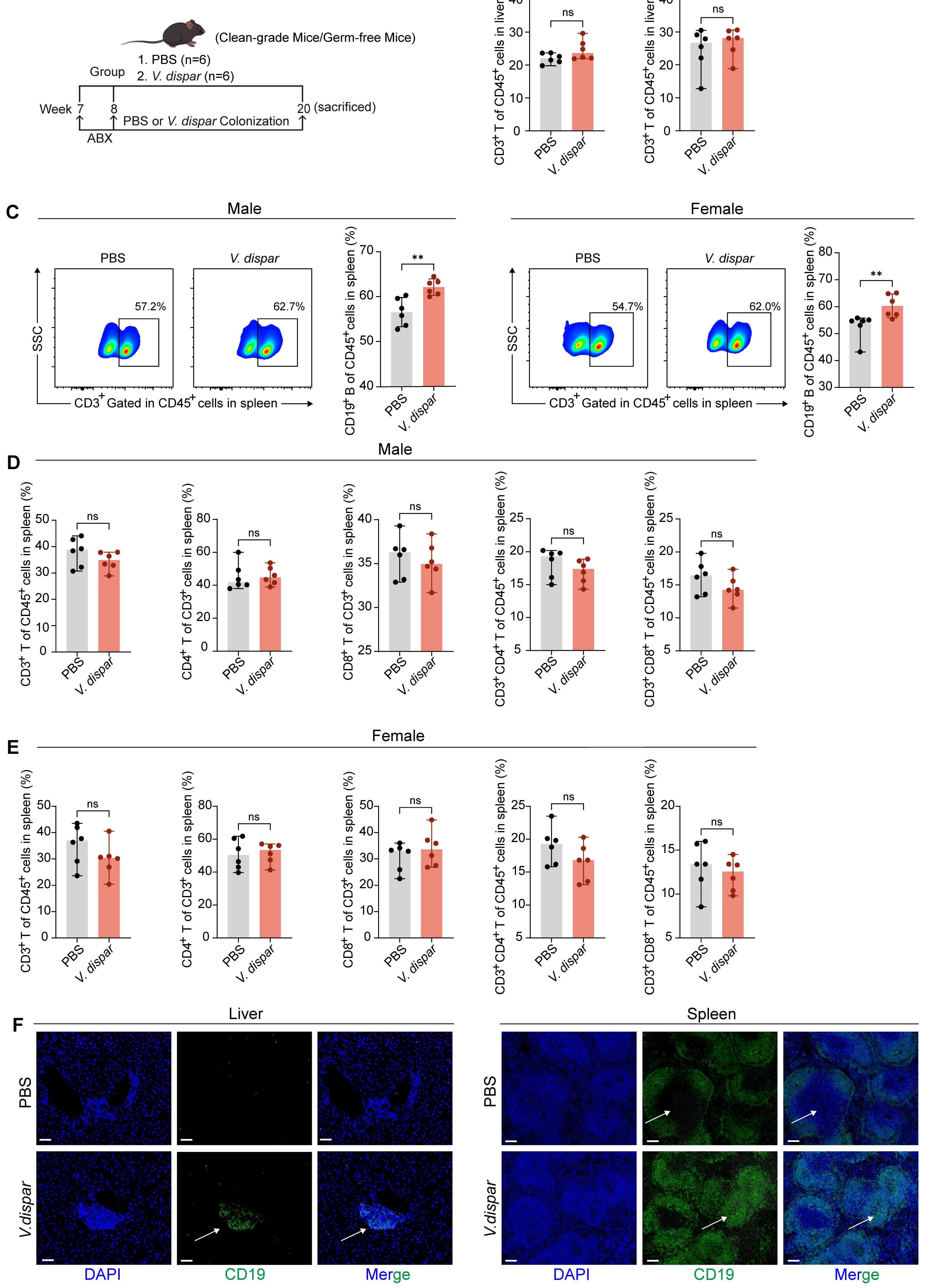
A
(Clean-grade Mice/Germ-free Mice)
Group
1. PBS (n=6)
2. V. dispar (n=6)
Week 7 8 20 (sacrificed)
PBS or V. dispar Colonization
ABX
B
Male
Female
CD3+ T of CD45+ cells in liver (%)
ns
PBS
V. dispar
C
Male
Female
PBS
V. dispar
57.2%
62.7%
54.7%
62.0%
SSC
CD3+ Gated in CD45+ cells in spleen
CD19+ B of CD45+ cells in spleen (%)
**
D
Male
CD3+ T of CD45+ cells in spleen (%)
CD4+ T of CD3+ cells in spleen (%)
CD8+ T of CD3+ cells in spleen (%)
CD3+CD4+ T of CD45+ cells in spleen (%)
CD3+CD8+ T of CD45+ cells in spleen (%)
E
Female
F
Liver
Spleen
PBS
V.dispar
DAPI
CD19
Merge

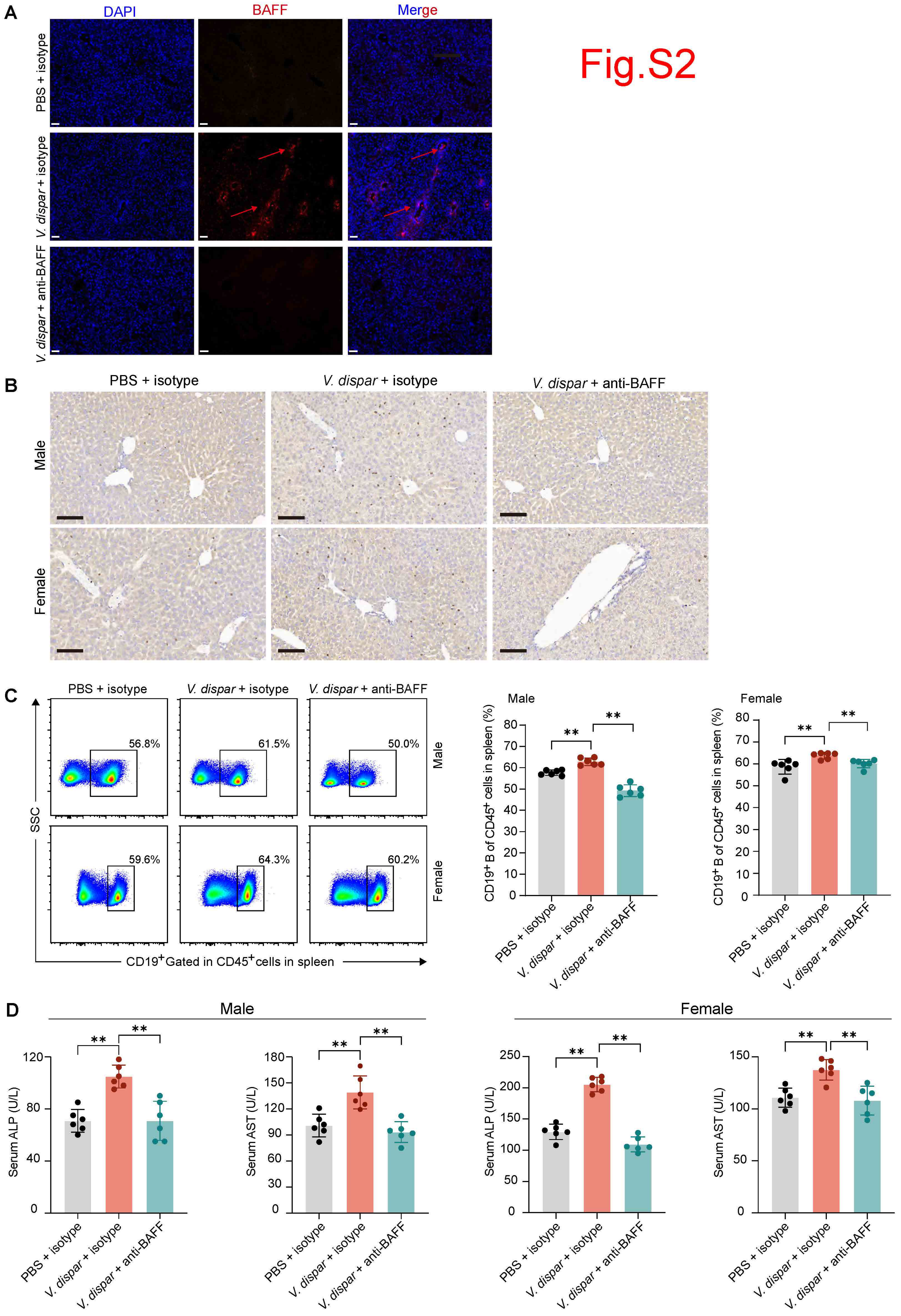
Fig.S2
A
DAPI
BAFF
Merge
PBS + isotype
V. dispar + isotype
V. dispar + anti-BAFF
B
PBS + isotype
V. dispar + isotype
V. dispar + anti-BAFF
Male
Female
C
PBS + isotype
V. dispar + isotype
V. dispar + anti-BAFF
56.8%
61.5%
50.0%
59.6%
64.3%
60.2%
Male
Female
SSC
CD19+ Gated in CD45+ cells in spleen
Male
Female
CD19+ B of CD45+ cells in spleen (%)
**
D
Male
Female
Serum ALP (U/L)
Serum AST (U/L)
**

Fig.S3

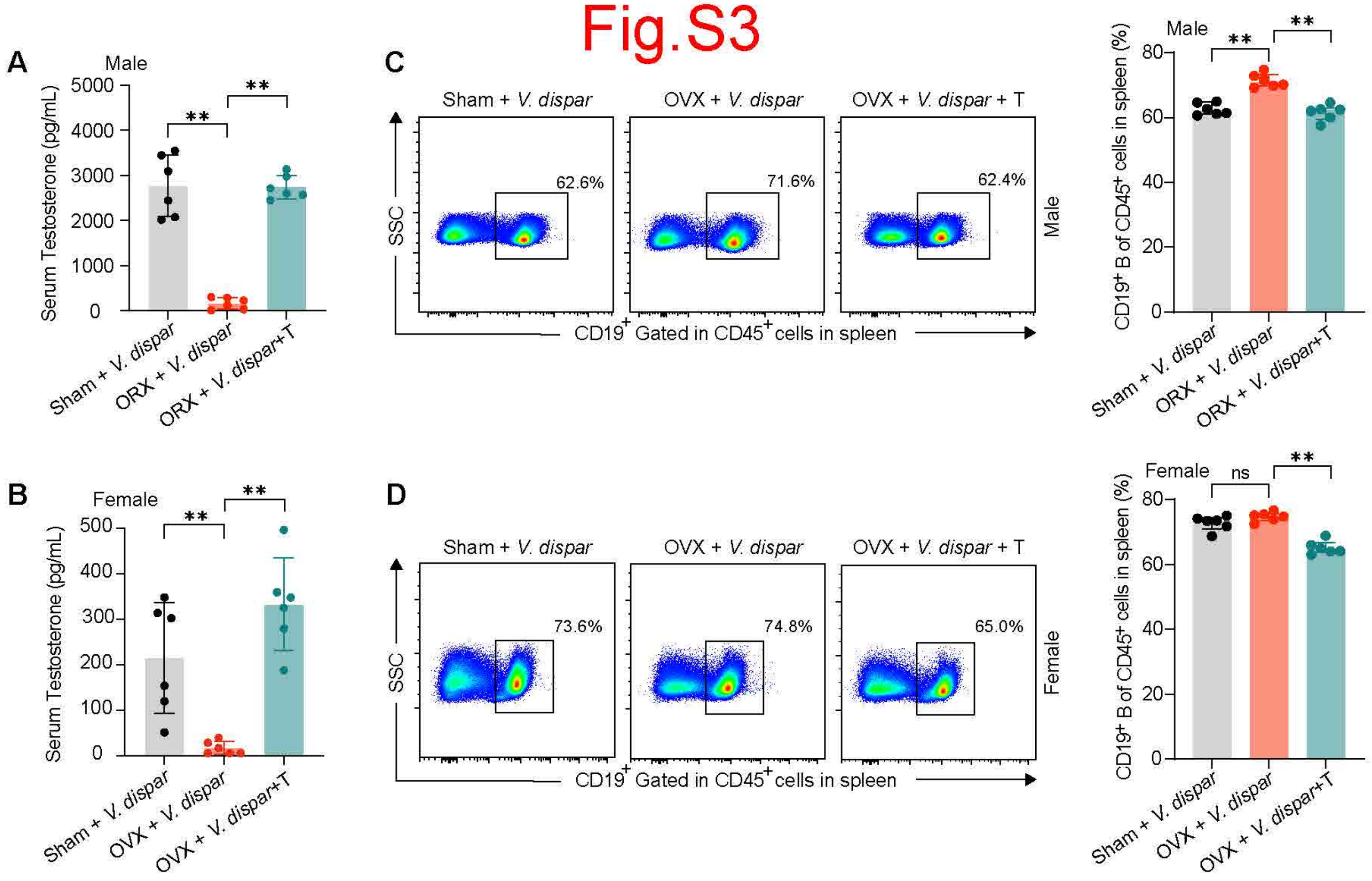

A
Male
Serum Testosterone (pg/mL)
5000
4000
3000
2000
1000
0
**
**
Sham + V. dispar
ORX + V. dispar
ORX + V. dispar+T
B
Female
Serum Testosterone (pg/mL)
500
400
300
200
100
0
**
**
Sham + V. dispar
OVX + V. dispar
OVX + V. dispar+T
C
Sham + V. dispar
OVX + V. dispar
OVX + V. dispar + T
SSC
62.6%
71.6%
62.4%
Male
CD19+ Gated in CD45+ cells in spleen
Male
CD19+ B of CD45+ cells in spleen (%)
80
60
40
20
0
**
**
Sham + V. dispar
ORX + V. dispar
ORX + V. dispar+T
D
Sham + V. dispar
OVX + V. dispar
OVX + V. dispar + T
SSC
73.6%
74.8%
65.0%
Female
CD19+ Gated in CD45+ cells in spleen
Female
CD19+ B of CD45+ cells in spleen (%)
80
60
40
20
0
ns
**
Sham + V. dispar
OVX + V. dispar
OVX + V. dispar+T

### Fig.S4

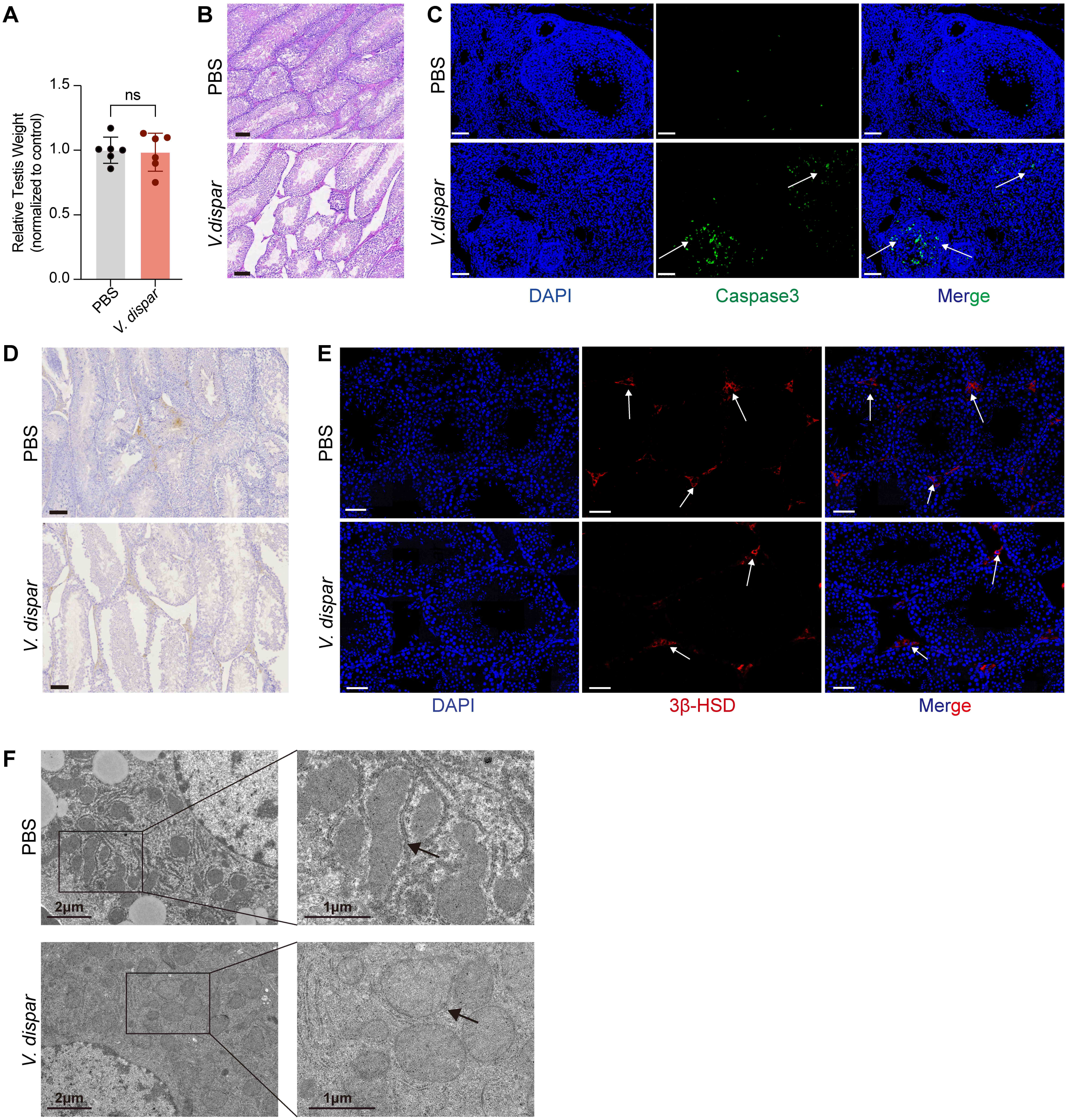
A
Relative Testis Weight (normalized to control)
1.5
1.0
0.5
0.0
ns
PBS
V. dispar
B
PBS
V.dispar
C
PBS
V.dispar
DAPI
Caspase3
Merge
D
PBS
V. dispar
E
PBS
V. dispar
DAPI
3β-HSD
Merge
F
PBS
V. dispar
2μm
1μm
2μm
1μm

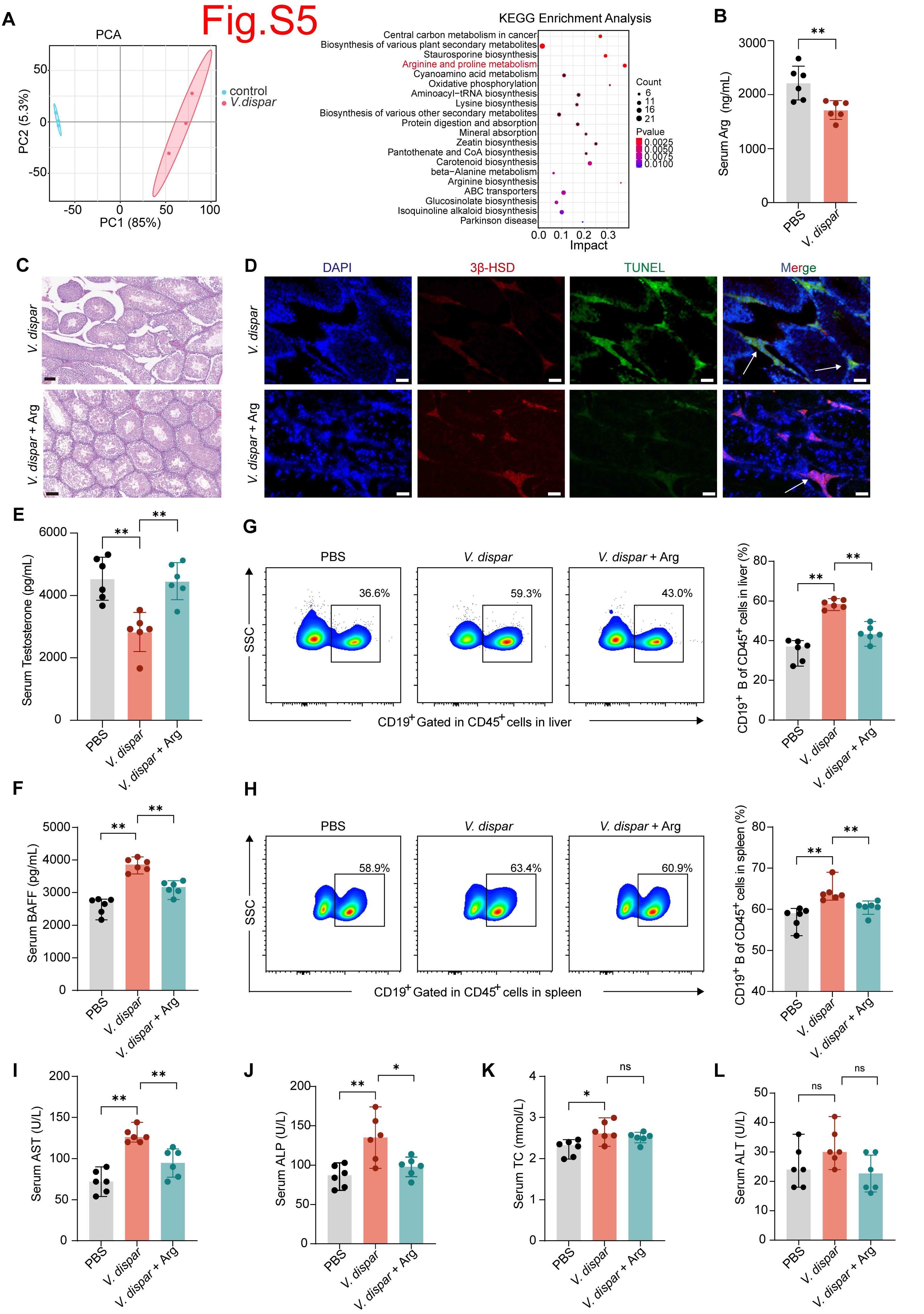
Fig.S5
A
PCA
control
V.dispar
PC1 (85%)
PC2 (5.3%)
KEGG Enrichment Analysis
Central carbon metabolism in cancer
Biosynthesis of various plant secondary metabolites
Staurosporine biosynthesis
Arginine and proline metabolism
Cyanoamino acid metabolism
Oxidative phosphorylation
Aminoacyl−tRNA biosynthesis
Lysine biosynthesis
Biosynthesis of various other secondary metabolites
Protein digestion and absorption
Mineral absorption
Zeatin biosynthesis
Pantothenate and CoA biosynthesis
Carotenoid biosynthesis
beta−Alanine metabolism
Arginine biosynthesis
ABC transporters
Glucosinolate biosynthesis
Isoquinoline alkaloid biosynthesis
Parkinson disease
Count
Pvalue
Impact
B
Serum Arg (ng/mL)
PBS
V. dispar
C
V. dispar
V. dispar + Arg
D
DAPI
3β-HSD
TUNEL
Merge
E
Serum Testosterone (pg/mL)
G
PBS
V. dispar
V. dispar + Arg
36.6%
59.3%
43.0%
SSC
CD19+ Gated in CD45+ cells in liver
CD19+ B of CD45+ cells in liver (%)
F
Serum BAFF (pg/mL)
H
58.9%
63.4%
60.9%
CD19+ Gated in CD45+ cells in spleen
CD19+ B of CD45+ cells in spleen (%)
I
Serum AST (U/L)
J
Serum ALP (U/L)
K
Serum TC (mmol/L)
L
Serum ALT (U/L)
ns

### Fig.S6

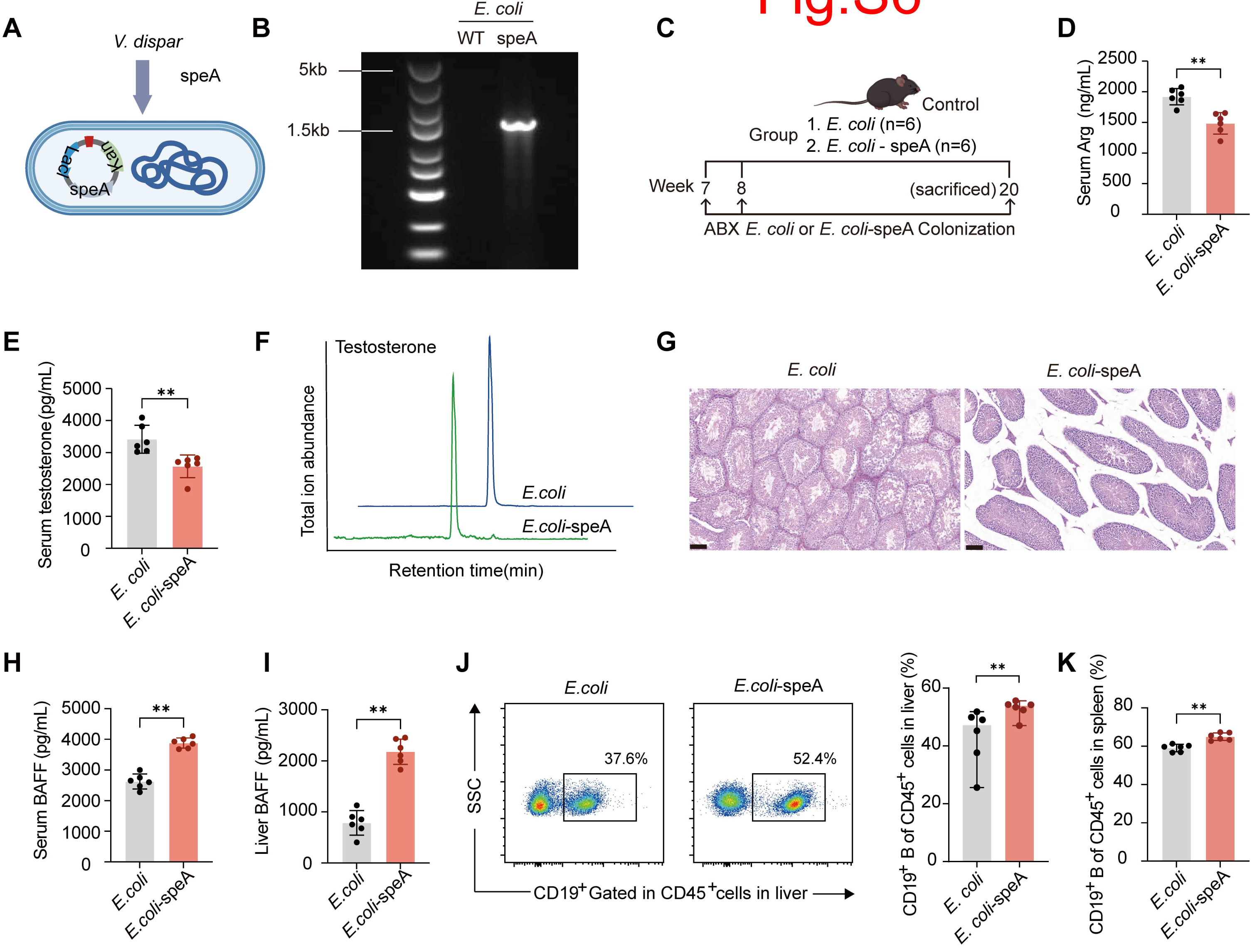

A
V. dispar
speA
LacI
Kan
speA
B
E. coli
WT speA
5kb
1.5kb
C
Control
Group
1. E. coli (n=6)
2. E. coli - speA (n=6)
Week 7 8 (sacrificed) 20
ABX E. coli or E. coli-speA Colonization
D
Serum Arg (ng/mL)
2500
2000
1500
1000
500
0
**
E. coli
E. coli-speA
E
Serum testosterone (pg/mL)
5000
4000
3000
2000
1000
0
**
E. coli
E. coli-speA
F
Testosterone
Total ion abundance
E.coli
E.coli-speA
Retention time(min)
G
E. coli
E. coli-speA
H
Serum BAFF (pg/mL)
5000
4000
3000
2000
1000
0
**
E.coli
E.coli-speA
I
Liver BAFF (pg/mL)
3000
2000
1000
0
**
E.coli
E.coli-speA
J
E.coli
E.coli-speA
37.6%
52.4%
SSC
CD19+ Gated in CD45+ cells in liver
CD19+ B of CD45+ cells in liver (%)
60
40
20
0
**
E. coli
E. coli-speA
K
CD19+ B of CD45+ cells in spleen (%)
80
60
40
20
0
**
E.coli
E.coli-speA

### Fig.S7

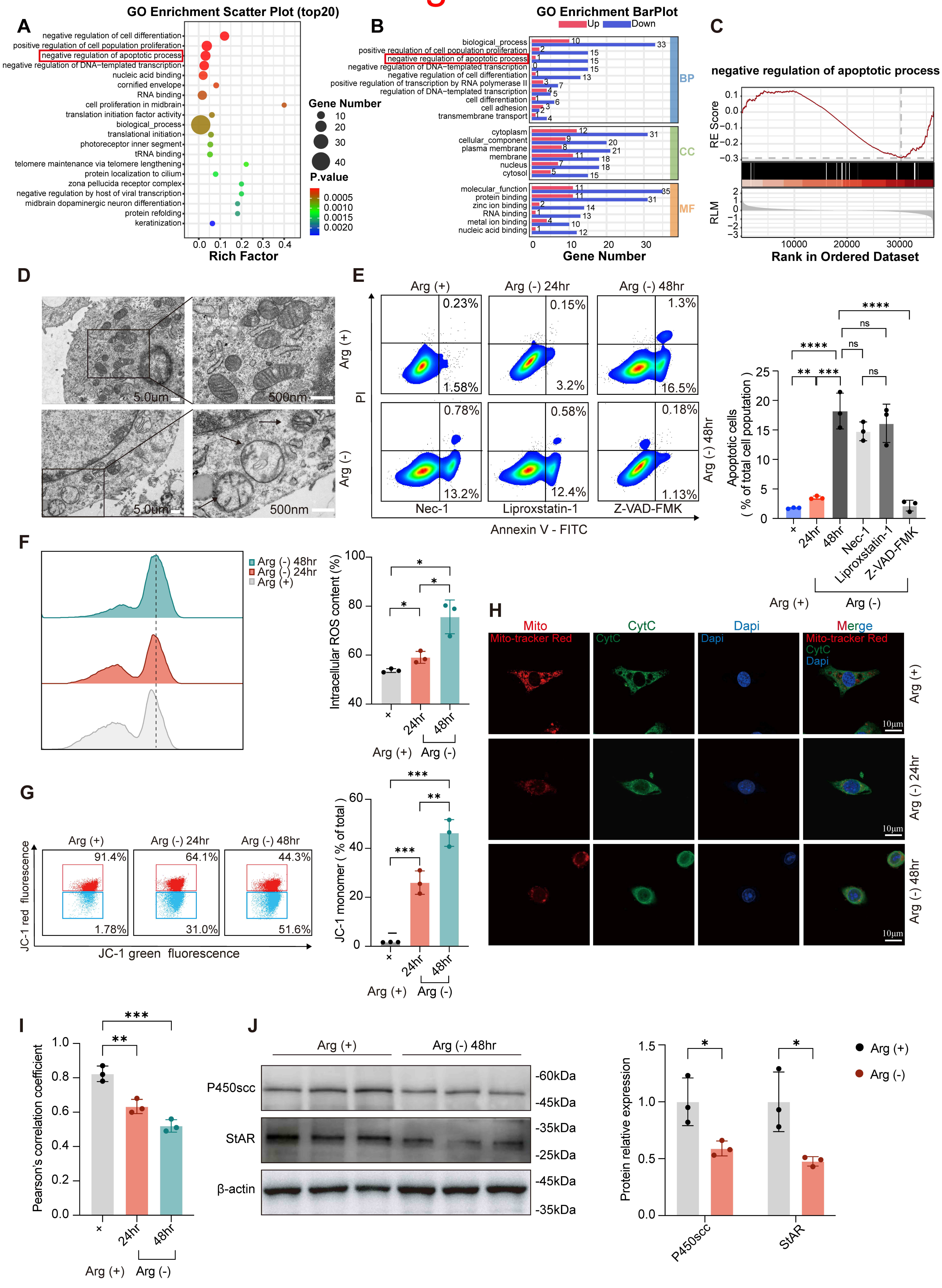
A
GO Enrichment Scatter Plot (top20)
negative regulation of cell differentiation
positive regulation of cell population proliferation
negative regulation of apoptotic process
negative regulation of DNA−templated transcription
nucleic acid binding
cornified envelope
RNA binding
cell proliferation in midbrain
translation initiation factor activity
biological_process
translational initiation
photoreceptor inner segment
tRNA binding
telomere maintenance via telomere lengthening
protein localization to cilium
zona pellucida receptor complex
negative regulation by host of viral transcription
midbrain dopaminergic neuron differentiation
protein refolding
keratinization
Rich Factor
Gene Number
P.value
B
GO Enrichment BarPlot
Up
Down
BP
CC
MF
Gene Number
C
negative regulation of apoptotic process
RE Score
RLM
Rank in Ordered Dataset
D
5.0um
500nm
E
Arg (+)
Arg (-) 24hr
Arg (-) 48hr
Nec-1
Liproxstatin-1
Z-VAD-FMK
PI
Annexin V - FITC
Apoptotic cells
( % of total cell popuation )
F
Arg (-) 48hr
Arg (-) 24hr
Arg (+)
Intracellular ROS content (%)
G
JC-1 red fluorescence
JC-1 green fluorescence
JC-1 monomer ( % of total )
H
Mito
CytC
Dapi
Merge
Mito-tracker Red
10μm
I
Pearson's correlation coefficient
J
P450scc
StAR
β-actin
-60kDa
-45kDa
-35kDa
-25kDa
Protein relative expression
Arg (+)
Arg (-)

Fig.S8

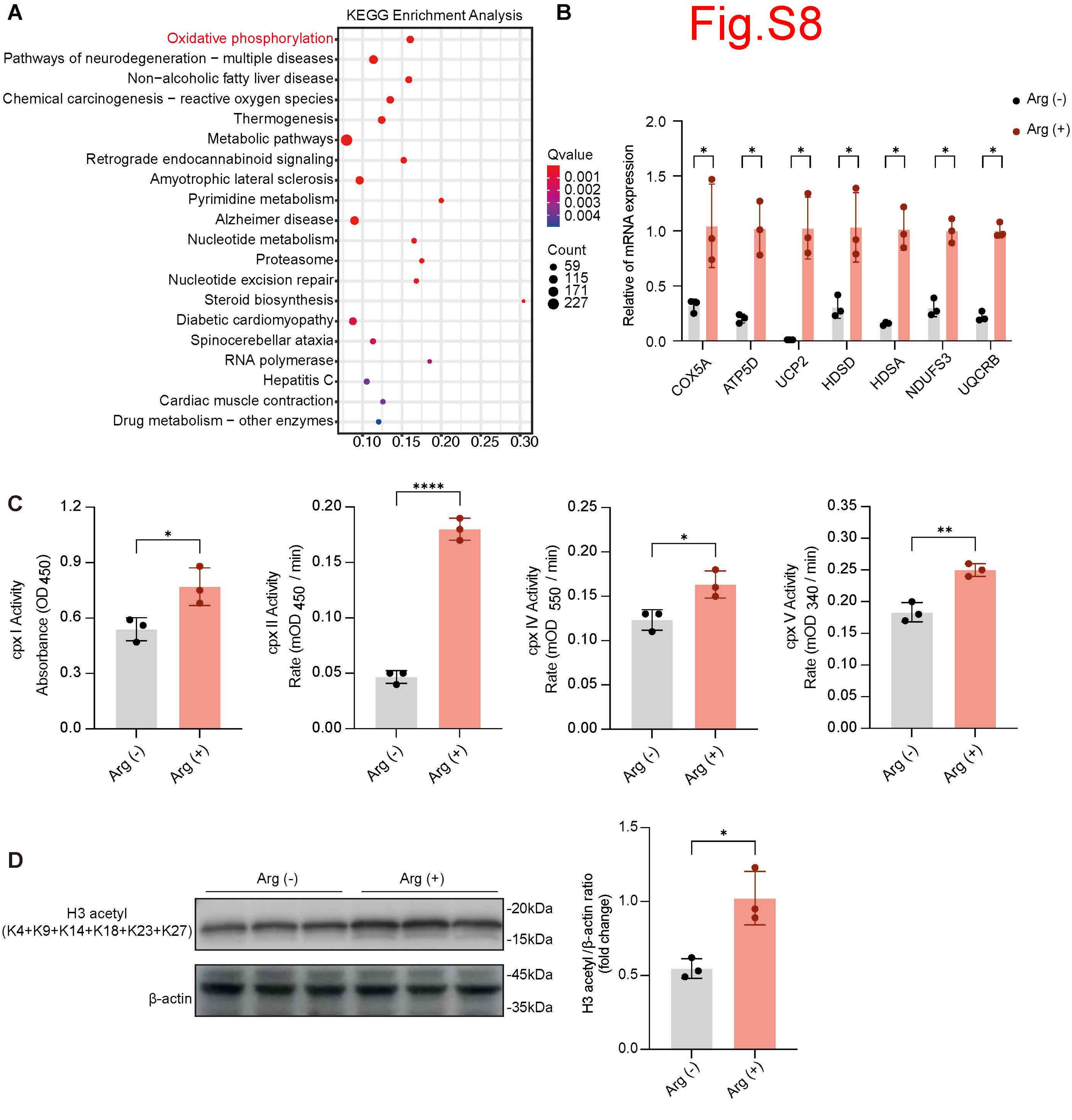
A
KEGG Enrichment Analysis
Oxidative phosphorylation
Pathways of neurodegeneration − multiple diseases
Non−alcoholic fatty liver disease
Chemical carcinogenesis − reactive oxygen species
Thermogenesis
Metabolic pathways
Retrograde endocannabinoid signaling
Amyotrophic lateral sclerosis
Pyrimidine metabolism
Alzheimer disease
Nucleotide metabolism
Proteasome
Nucleotide excision repair
Steroid biosynthesis
Diabetic cardiomyopathy
Spinocerebellar ataxia
RNA polymerase
Hepatitis C
Cardiac muscle contraction
Drug metabolism − other enzymes
0.10 0.15 0.20 0.25 0.30
Qvalue
0.001
0.002
0.003
0.004
Count
59
115
171
227
B
Arg (-)
Arg (+)
Relative of mRNA expression
2.0
1.5
1.0
0.5
0.0
COX5A
ATP5D
UCP2
HDSD
HDSA
NDUFS3
UQCRB
C
cpx I Activity
Absorbance (OD 450)
cpx II Activity
Rate (mOD 450 / min)
cpx IV Activity
Rate (mOD 550 / min)
cpx V Activity
Rate (mOD 340 / min)
Arg (-)
Arg (+)
D
Arg (-)
Arg (+)
H3 acetyl
(K4+K9+K14+K18+K23+K27)
β-actin
-20kDa
-15kDa
-45kDa
-35kDa
H3 acetyl /β-actin ratio
(fold change)

Fig.S9-1

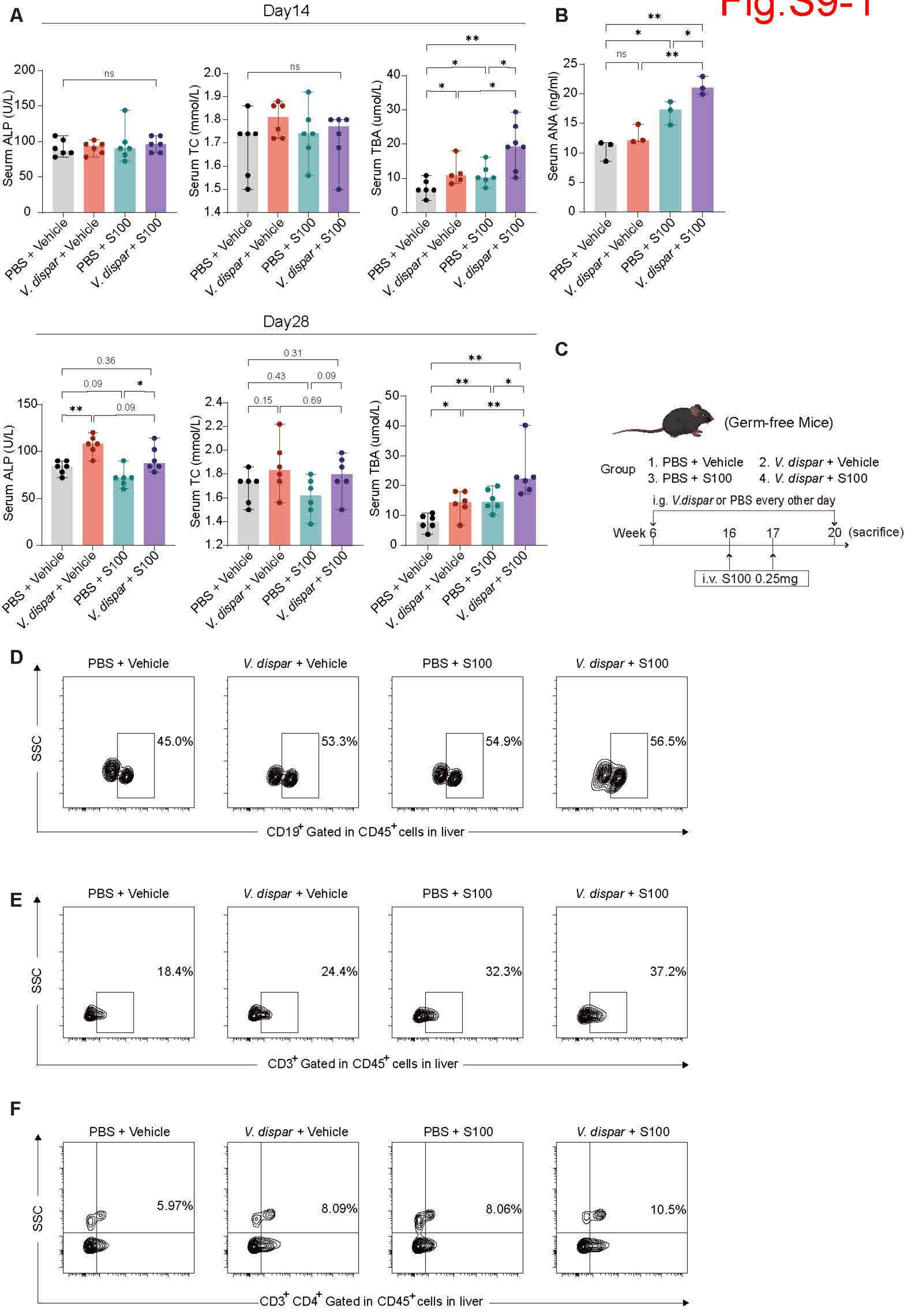

A
Day14
Seurm ALP (U/L)
Serum TC (mmol/L)
Serum TBA (umol/L)
ns
PBS + Vehicle
V. dispar + Vehicle
PBS + S100
V. dispar + S100
B
Serum ANA (ng/ml)
Day28
0.36
0.09
0.31
0.43
0.15
0.69
Serum ALP (U/L)
C
(Germ-free Mice)
Group
1. PBS + Vehicle
2. V. dispar + Vehicle
3. PBS + S100
4. V. dispar + S100
i.g. V.dispar or PBS every other day
Week 6
16
17
20
(sacrifice)
i.v. S100 0.25mg
D
SSC
45.0%
53.3%
54.9%
56.5%
CD19+ Gated in CD45+ cells in liver
E
18.4%
24.4%
32.3%
37.2%
CD3+ Gated in CD45+ cells in liver
F
5.97%
8.09%
8.06%
10.5%
CD3+ CD4+ Gated in CD45+ cells in liver

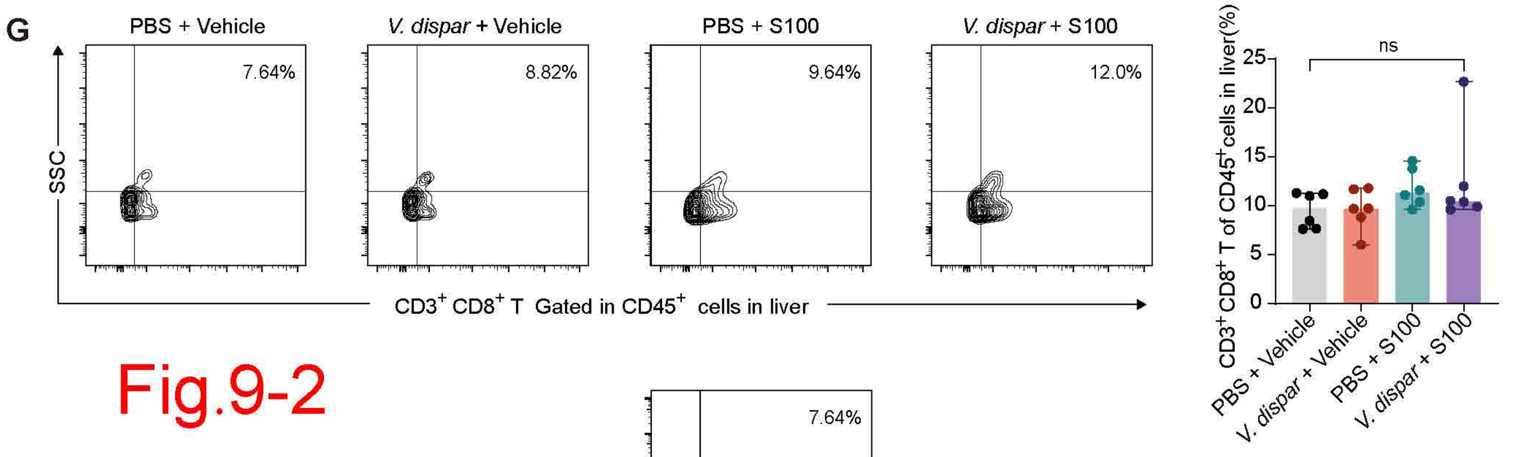
G
PBS + Vehicle
V. dispar + Vehicle
PBS + S100
V. dispar + S100
7.64%
8.82%
9.64%
12.0%
SSC
CD3+ CD8+ T Gated in CD45+ cells in liver
ns
CD3+ CD8+ T of CD45+ cells in liver(%)
25
20
15
10
5
0
PBS + Vehicle
V. dispar + Vehicle
PBS + S100
V. dispar + S100

Fig.9-2

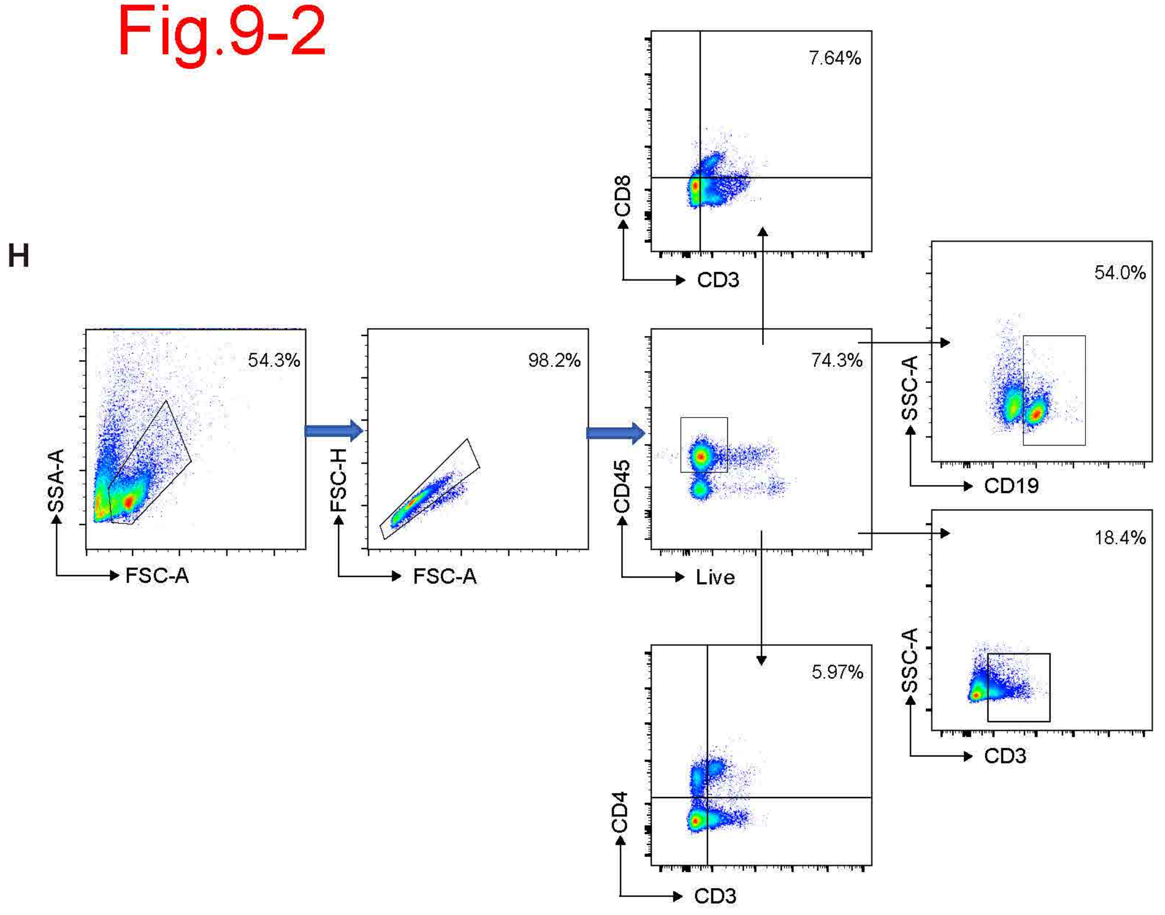
H
7.64%
CD8
CD3
54.0%
SSC-A
CD19
54.3%
SSA-A
FSC-A
98.2%
FSC-H
FSC-A
74.3%
CD45
Live
18.4%
SSC-A
CD3
5.97%
CD4
CD3

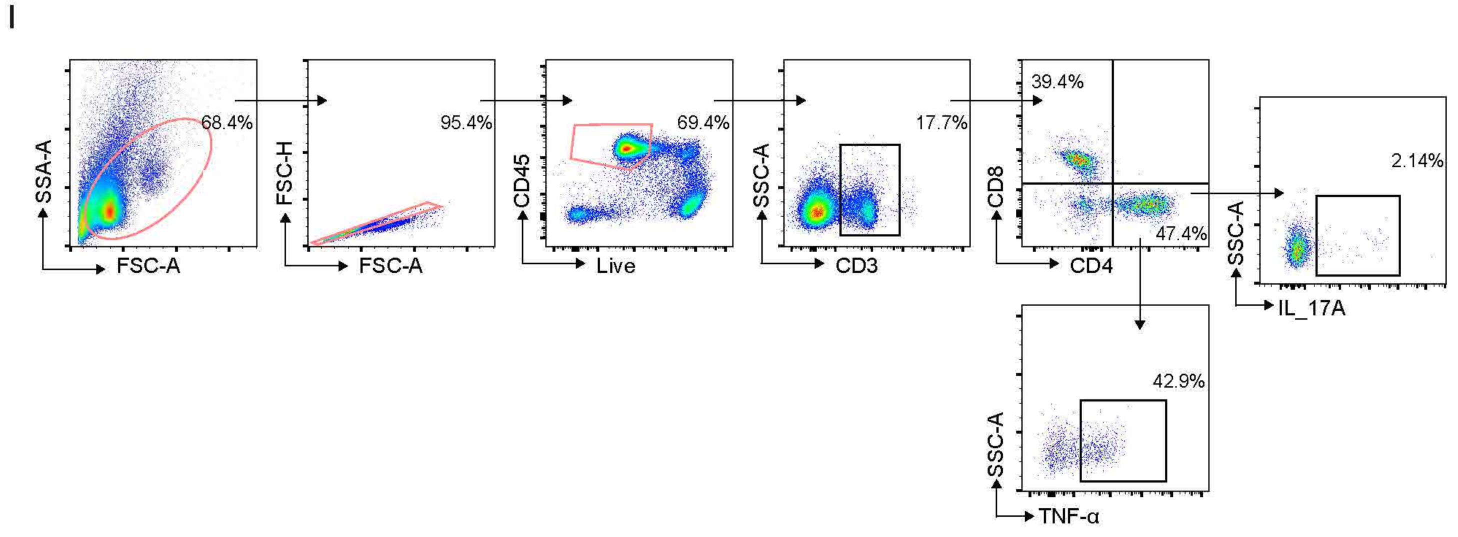
I
68.4%
SSA-A
FSC-A
95.4%
FSC-H
FSC-A
69.4%
CD45
Live
17.7%
SSC-A
CD3
39.4%
47.4%
CD8
CD4
2.14%
SSC-A
IL_17A
42.9%
SSC-A
TNF-α

Fig.S10

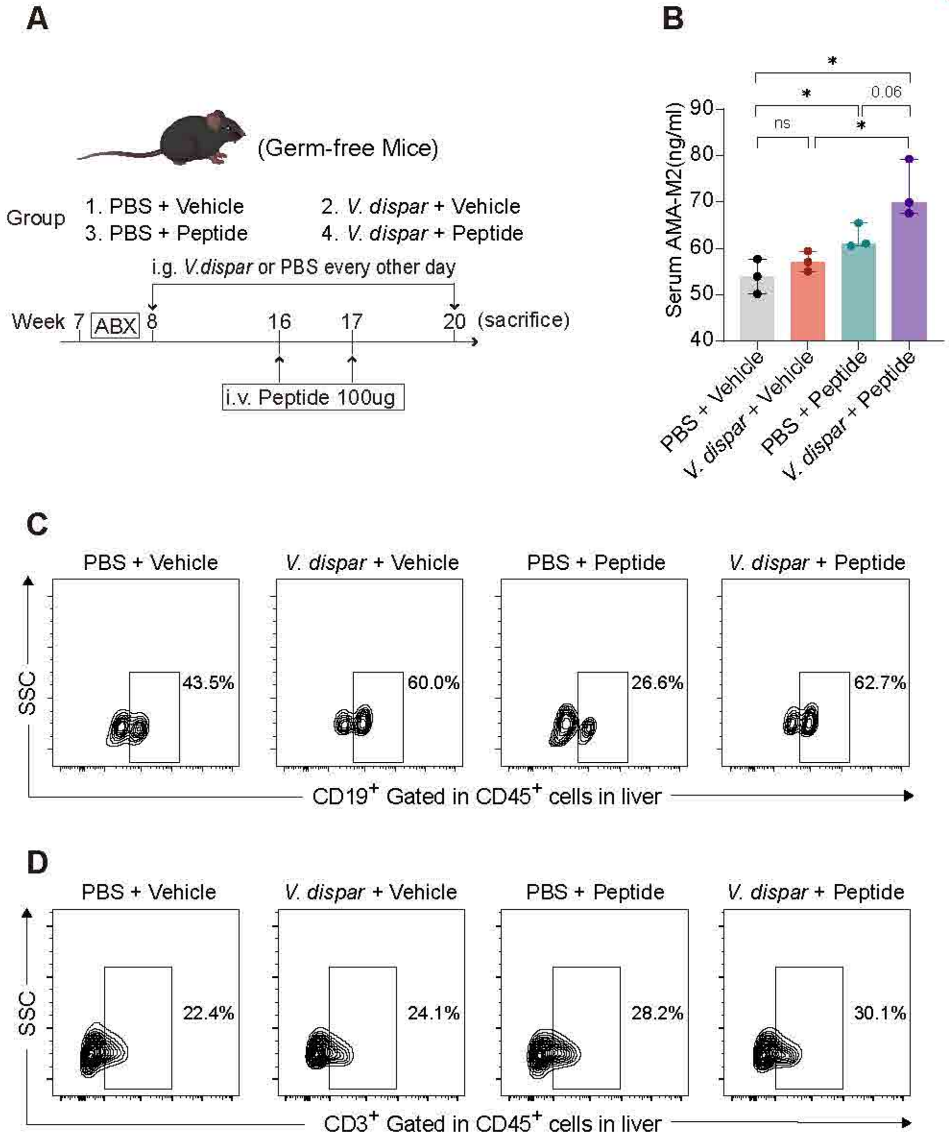

A
(Germ-free Mice)
Group
1. PBS + Vehicle
2. V. dispar + Vehicle
3. PBS + Peptide
4. V. dispar + Peptide
i.g. V.dispar or PBS every other day
Week 7
ABX
8
16
17
20
(sacrifice)
i.v. Peptide 100ug
B
Serum AMA-M2(ng/ml)
ns
*
0.06
PBS + Vehicle
V. dispar + Vehicle
PBS + Peptide
V. dispar + Peptide
C
PBS + Vehicle
V. dispar + Vehicle
PBS + Peptide
V. dispar + Peptide
SSC
43.5%
60.0%
26.6%
62.7%
CD19+ Gated in CD45+ cells in liver
D
PBS + Vehicle
V. dispar + Vehicle
PBS + Peptide
V. dispar + Peptide
SSC
22.4%
24.1%
28.2%
30.1%
CD3+ Gated in CD45+ cells in liver
